# High quality Bathyarchaeia MAGs from lignocellulose-impacted environments elucidate metabolism and evolutionary mechanisms

**DOI:** 10.1101/2024.09.16.613360

**Authors:** Camilla Lothe Nesbø, Ilya Kublanov, Minqing Yang, Anupama Achal Sharan, Torsten Meyer, Elizabeth A. Edwards

**Author notes:** **Competing Interests**. The authors declare no competing financial interests.

## Abstract

The archaeal class Bathyarchaeia are widely and abundantly distributed in anoxic habitats. Metagenomic studies have suggested that they are mixotrophic, capable of CO_2_ fixation and heterotrophic growth, and involved in acetogenesis and lignin degradation. We analysed 35 Bathyarchaeia metagenome assembled genomes (MAGs), including the first complete circularized MAG (cMAG) of the Bathy-6 subgroup, from the metagenomes of three full scale pulp and paper mill anaerobic digesters and three laboratory methanogenic enrichment cultures maintained on pre-treated poplar. Thirty-three MAGs belong to the Bathy-6 lineage while two are from the Bathy-8 lineage. In our previous analysis of the microbial community in the pulp mill digesters, Bathyarchaeia were abundant and positively correlated to hydrogenotrophic and methylotrophic methanogenesis. Several factors likely contribute to the success of the Bathy-6 lineage compared to Bathy-8 in the reactors. The Bathy-6 genomes are larger than those of Bathy-8 and have more genes involved in lignocellulose degradation including carbohydrate-active enzymes (CAZymes) not present in the Bathy-8. Bathy-6 also share the Bathyarchaeal *O*-demethylase-system recently identified in Bathy-8. All the Bathy-6 MAGs had numerous membrane-associated pyrroloquinoline quinone-domain (PQQ) proteins that we suggest are involved in lignin modification or degradation, together with Radical-SAM and Rieske domain proteins, and AA2, AA3, and AA6-family oxidoreductases. We also identified a complete B12 synthesis pathway and a complete nitrogenase gene locus. Finally, comparative genomic analyses revealed that Bathyarchaeia genomes are dynamic and have interacted with other organisms in their environments through gene transfer to expand their gene repertoire.

## INTRODUCTION

Bathyarchaeia, previously known as the Miscellaneous Crenarchaeota Group, are an archaeal class abundant in anoxic environments [1, 2]. They have been suggested to constitute one of the most abundant cell lineages on our planet and play an important role in global biogeochemical cycling [1, 2]. Although they are at low numbers in the anoxic human gut [3], Bathyarchaeia have been detected at high abundance in animals that enjoy lignocellulosic diets such as termites [4], beavers and moose [5].

Within the class Bathyarchaeia, twenty-five monophyletic subgroups (Bathy-1 - Bathy-25) were originally defined based on 16S rRNA gene phylogenies [1, 6]. Hou *et al*. [2] recently reclassified these into eight order level lineages. However, this proposed taxonomy is yet to be formally accepted. For instance, the 16S rRNA lineage we report on, Bathy-6, was assigned to order Baizomonadales and to genus *Candidatus* Baizomonas by Hou *et al*. [2] while Loh *et al*. [4] proposed two distinct genera *within* Bathy-6. In the genome taxonomy database (GTDB) v 214 [7] the lineages still have ‘place-holder’ names (Table S1B). Here we therefore refer to the lineages using the 16S-lineage designations (i.e. Bathy-6 and Bathy-8) and await an approved taxonomy.

Analyses of the functional potential of Bathyarchaeia metagenome assembled genomes (MAGs) suggest they can grow on a range of organic substrates [1, 2]. Although no pure culture of Bathyarcheia is available, two cultures highly enriched for organisms from the Bathy-8 lineage were recently found to grow on alkali lignin or the lignin degradation product, 3,4-dimethoxybenzoic acid [8, 9]. One of the cultures was shown to grow on proteinaceous substrates, but not carbohydrates [8]. A Bathy-8 population, previously shown to be stimulated by lignin [10], were recently reported to encode an *O*-demethylase complex that demethylates products of lignin degradation [9]. Lin *et al*. [11] found that the Bathy-6 lineage was enriched in cultures grown on the lignin degradation product syringaldehyde, while Yu *et al*. [9] identified the genes for the *O*-demethylase-protein complex also in Bathy-6 MAGs. Moreover, inverse stable isotope labelling suggested that Bathy-8 organisms incorporated both organic and inorganic carbon when incubated with lignin [12]. Thus, both Bathy-6 and Bathy-8 organisms are likely involved in anaerobic lignin degradation.

We investigated the microbial communities in anaerobic wastewater treatment digestors at three Canadian pulp and paper mills (MillA, MillB and MillC) where samples were collected approximately twice a month over 1.5 years. Mill operational data and microbial community profiles were analyzed in Meyer *et al*. [13]. Bathyarchaeia 16S rRNA gene amplicon sequence variants (16S ASVs) were identified as part of the core community despite very different environmental conditions at each mill and can represent up to 19% of the archaeal community [13]. In MillC they positively correlated with hydrogenotrophic and methylotrophic methanogens and were, together with Dysgonomonadaceae, the only lineages linked to the colour concentration in the final wastewater after aerobic treatment [13], where colour formation results from oxidation of lignin degradation products [14]. This suggested a role of Bathyarchaeia in lignin degradation also in the pulp mill waste. Metagenome sequencing (Table S1) allowed us to assemble 16 MAGs classified as Bathyarchaeia, one of which represents the first closed circular genome Bathy-6 genome (cMAG). In addition, we assembled 19 Bathyarchaeia MAGs from three previously published metagenomes obtained from enrichment cultures amended with poplar hydrolysate [5, 15] (Table S1A). One of these enrichment cultures was created by inoculation with anaerobic granules from MillA. The two other cultures were seeded with beaver droppings and moose rumen, both environments expected to harbour microorganisms capable of lignocellulose degradation.

Here we analyse the genomic composition of our cMAG and compare it to the other MAGs to gain insights into the ecological role of Bathy-6 in lignocellulose-rich environments. The closed genome helped anchor the analyses and improved predictions of metabolism and evolutionary mechanisms.

## MATERIAL AND METHODS

### Metagenome Sources and Metagenome Assembled Genomes (MAGs)

The three mills investigated are described in detail in Meyer *et al*. [13]. Briefly, Mill A combines a chemical pulp mill and a bleached chemical thermo-mechanical pulp (BCTMP) mill, and Mills B and C are both BCTMP mills. Mill A has two internal circulation reactors, Mill B operates three anaerobic hybrid digesters, and Mill C has an anaerobic lagoon. Samples, DNA extraction, and sequencing of the samples from the pulp and paper mills are described in Meyer *et al*. [13], Nesbø *et al*. [16] and in Supplementary text S1. The enrichment cultures and their metagenomes are described in Wong *et al*. [5, 15]. Assembly and binning of the metagenomes from the mills and the enrichment cultures are described in Nesbø *et al*. [16] and in Supplementary text S1.

Taxonomy was assessed using the GTDB and GTDB-tk V.2.1.1 [17], and Bathyarchaeia MAGs were selected for further analyses. Completeness and redundancy values for the MAGs were obtained using CheckM [18]. The MAGs were further curated and refined as described in Supplementary text S2.

The MAGs were annotated using MetaErg [19]. Additional manual annotation was performed as described in Supplementary Text S2. Genomic loci were compared using Clinker [20]. Genome comparisons of high-quality MAGs to the cMAG were visualized using mummer2circos (https://github.com/metagenlab/mummer2circos).

### Phylogenomic, pangenome and phylogenetic analyses

Phylogenomic analysis of the MAGs was performed using the Gtotree-pipeline with the Archaea.hmm profile [21], including Bathyarchaeia-representative-genome sequences obtained from GTDB. The resulting alignment was imported to Geneious Prime v. 2022.0.1, sites with > 50% gaps were deleted, and a maximum likelihood tree was constructed using RAxML [22] with the WAG + GAMMA substitution model and 100 bootstrap replicates.

Pangenome analysis of the MAGs used the Anvi’o v. 7.1 [23] pangenome-snakemake-pipeline including closely related Bathyarchaeia genomes from the GTDB (Table S1C). Min-occurrence and MCL-inflation were set to 2. Calculation of average nucleotide identity (ANI) and annotation of the pangenome using the KEGG and the COG databases were also performed in Anvi’o v. 7.1.

Phylogenetic analyses of individual genes were performed in Geneious Prime v. 2022.0.1. Gene clusters (GCs) were extracted from the anvi’o pangenome, *in-silico* translated and compared against Genbank *nr* using BLASTP [24]. The top 100 matches were retrieved, aligned using MAFFT [25] and phylogenetic trees constructed using FastTree [26]. Default settings were used for all software.

## RESULTS

### Thirty-five Bathyarchaeia MAGs including the first closed Bathy-6 genome

Sixteen Bathyarchaeia MAGs were assembled from the six pulp mill metagenomes (Table 1), 10 of which contained 16S rRNA gene sequences identical to the ASVs in the time series amplicon data [13] (Fig. S1). The abundances measured by 16S ASVs and the metagenomic abundance data were similar (Table 1). We also assembled 19 Bathyarchaeia MAGs (Table 1) from metagenomes from enrichment cultures amended with either cellulose or poplar hydrolysate [5, 15]. Interestingly we found that Bathyarchaeia MAGs were observed at relatively high abundance in the poplar hydrolysate enrichments while they were at low or undetectable numbers in parallel enrichments on cellulose (Avicel) as sole carbon source (Table 1, Table S1E).

**Table 1.**
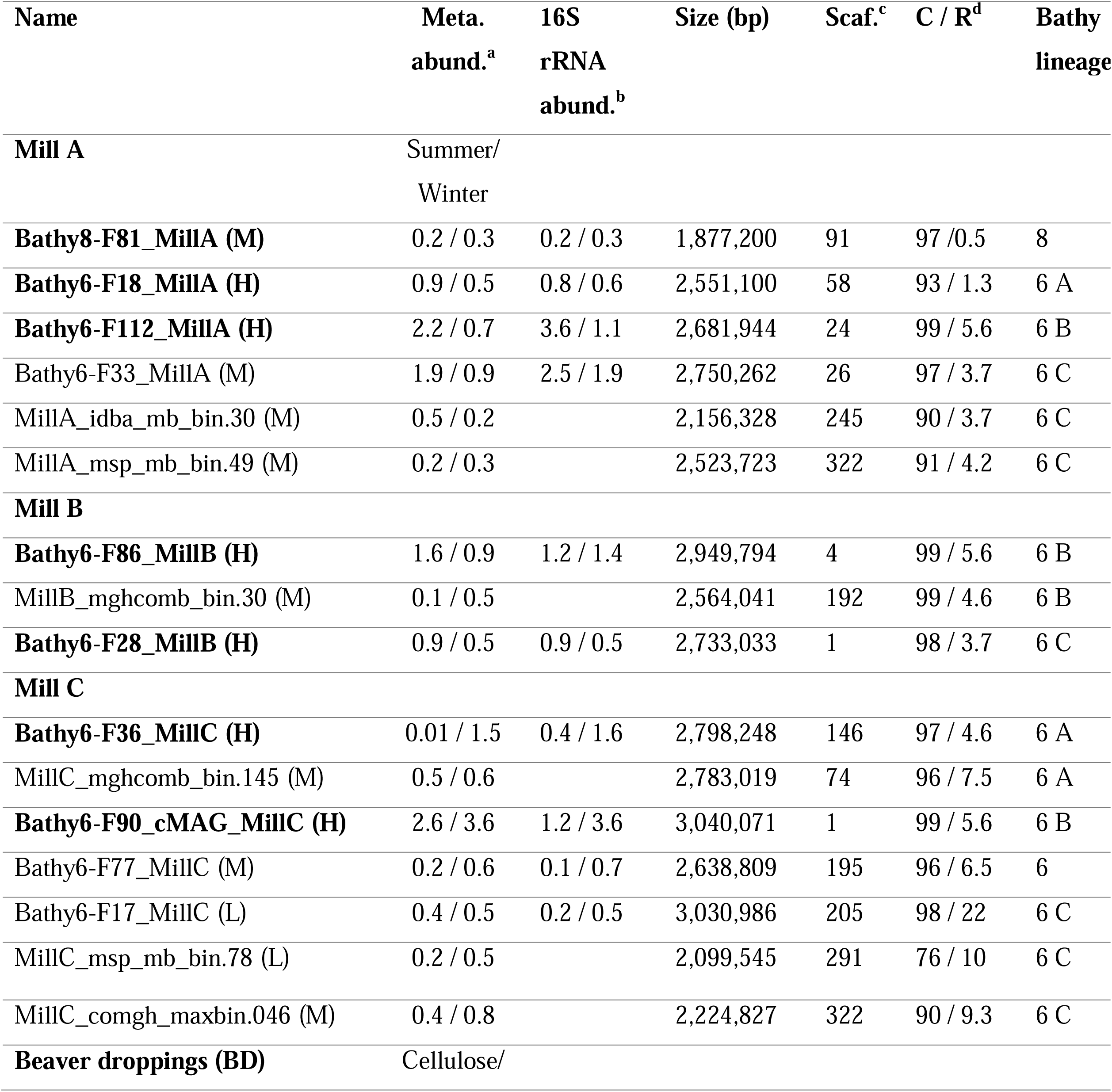
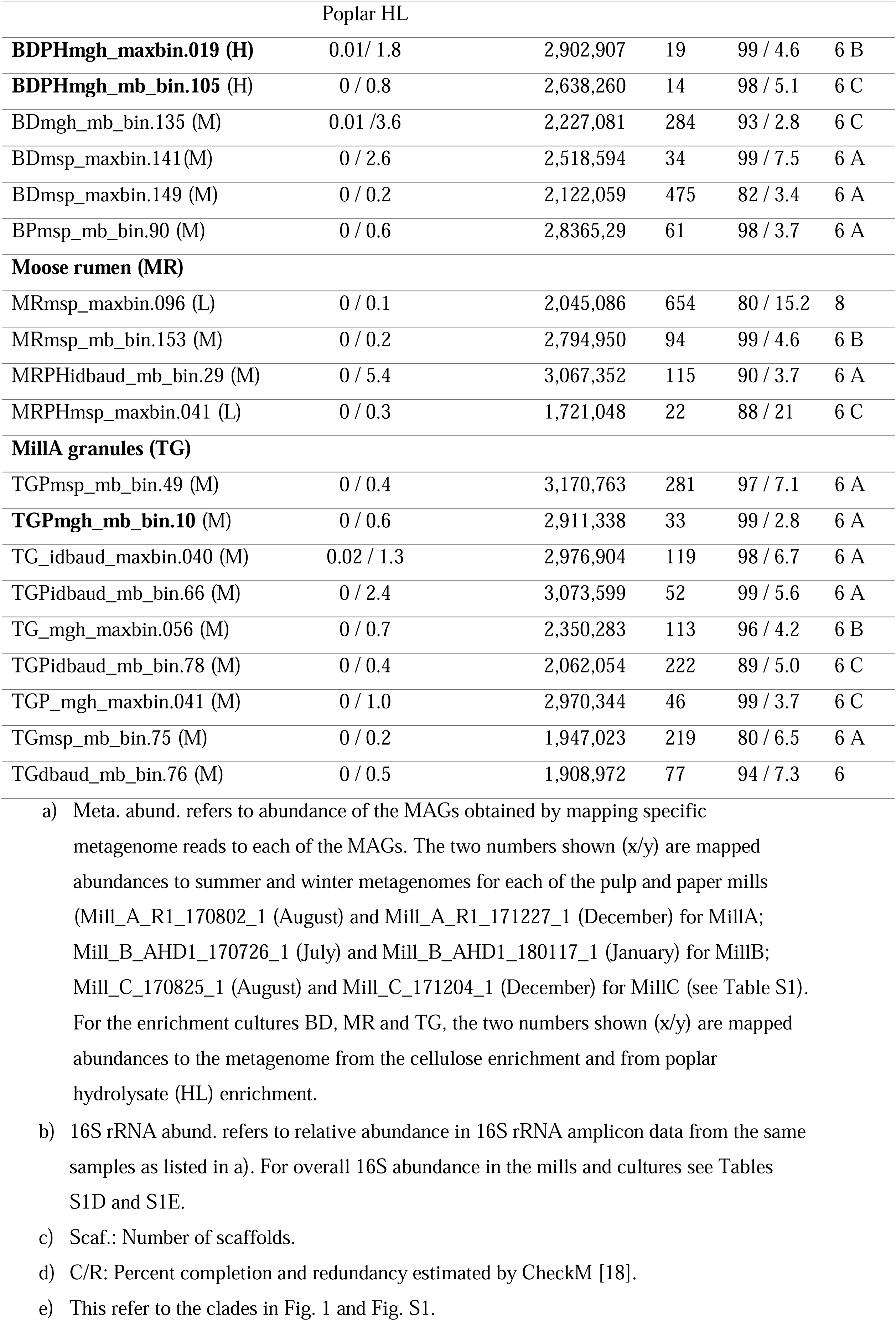
Overview of all Bathyarcheiea MAGs assembled by us. Name of the MAG indicates Bathyarchaeia lineage (Bathy-6 or Bathy-8), ASV feature id if 16S rRNA was identified (Fxx), and Mill the MAG originated from. Quality of the MAG is indicated in parenthesis H = high quality, M = medium quality and L = low quality following [29]. MAGs in bold were used in genome alignment in Fig. S7 and Fig. S11.

Most of the MAGs (33/35) clustered within the Bathy-6 clade in phylogenetic analyses of concatenated conserved single-copy protein sequences (Fig. 1) and 16S rRNA genes (Fig. S1).

**Figure 1.**
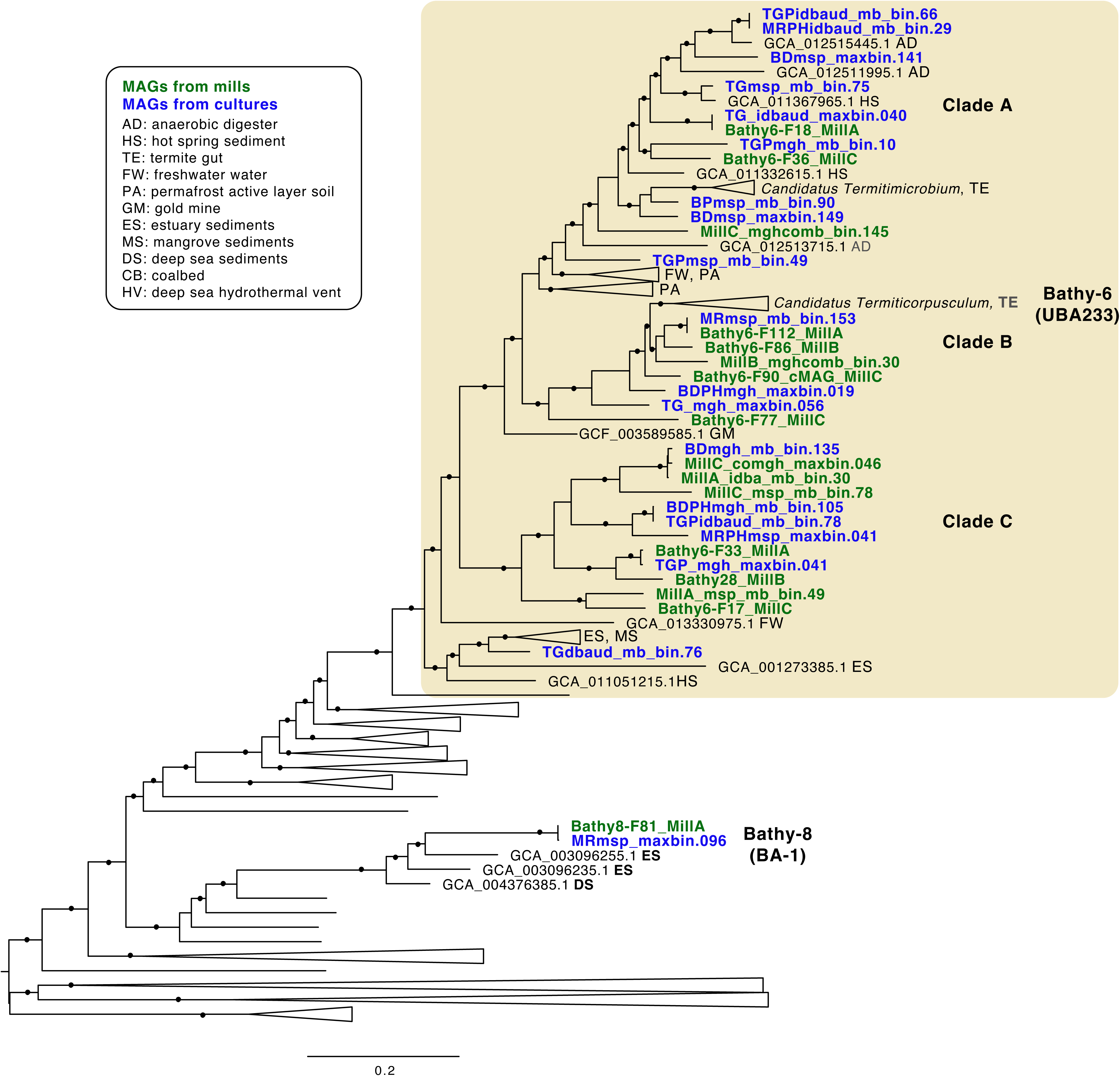
Maximum likelihood phylogenetic tree of Bathyarchaeia MAG and representative genomes from the genome taxonomy database. The tree is based on a 12,786 amino acid alignment of conserved single copy genes, generated by the gtotree pipeline. The tree was reconstructed using RAxML as implemented in Genenious Prime with the GAMMA WAG model and 100 bootstrap replicates. Black circles branches indicate bootstrap support values >= 70%. MAGs from the three paper mill digesters are given in green and MAGs from the three poplar hydrolysate enrichment cultures are in blue. Isolation sources have been added to representative MAGs from GTDB that were closely related to those generated in this study. Monophyletic clades with no close relatives in our MAG collection are collapsed and accession numbers associated with representative MAGs from GTDB not closely related to the ones generated here are not shown.

Two of the MAGs were assigned to Bathy-8. Average nucleotide identity (ANI) among all the MAGs suggested six species-level clusters [27] with ANI > =95% (Table S2). One MAG (MRmsp_maxbin_096) was a chimera of sequences from Bathy-8 and Bathy-6 with high ANI in comparison to both lineages (Fig. 2, Table S2) and was therefore excluded from gene content analyses. ANI within each of the Bathy-6 subclades (A, B and C in Fig. 1, Fig. 2) was 75-76%, while ANI between the clades was 70-72% (Table S2). We therefore propose that clade A, B and C represent different genera [28], and that clade A should be included in the proposed genus *Candidatus* Termitimicrobium [4] and clade B included in *Candidatus* Termiticorpusculum [4]. For clade C we propose the name *Candidatu*s Lignumamantes (wood lovers).

**Figure 2.**
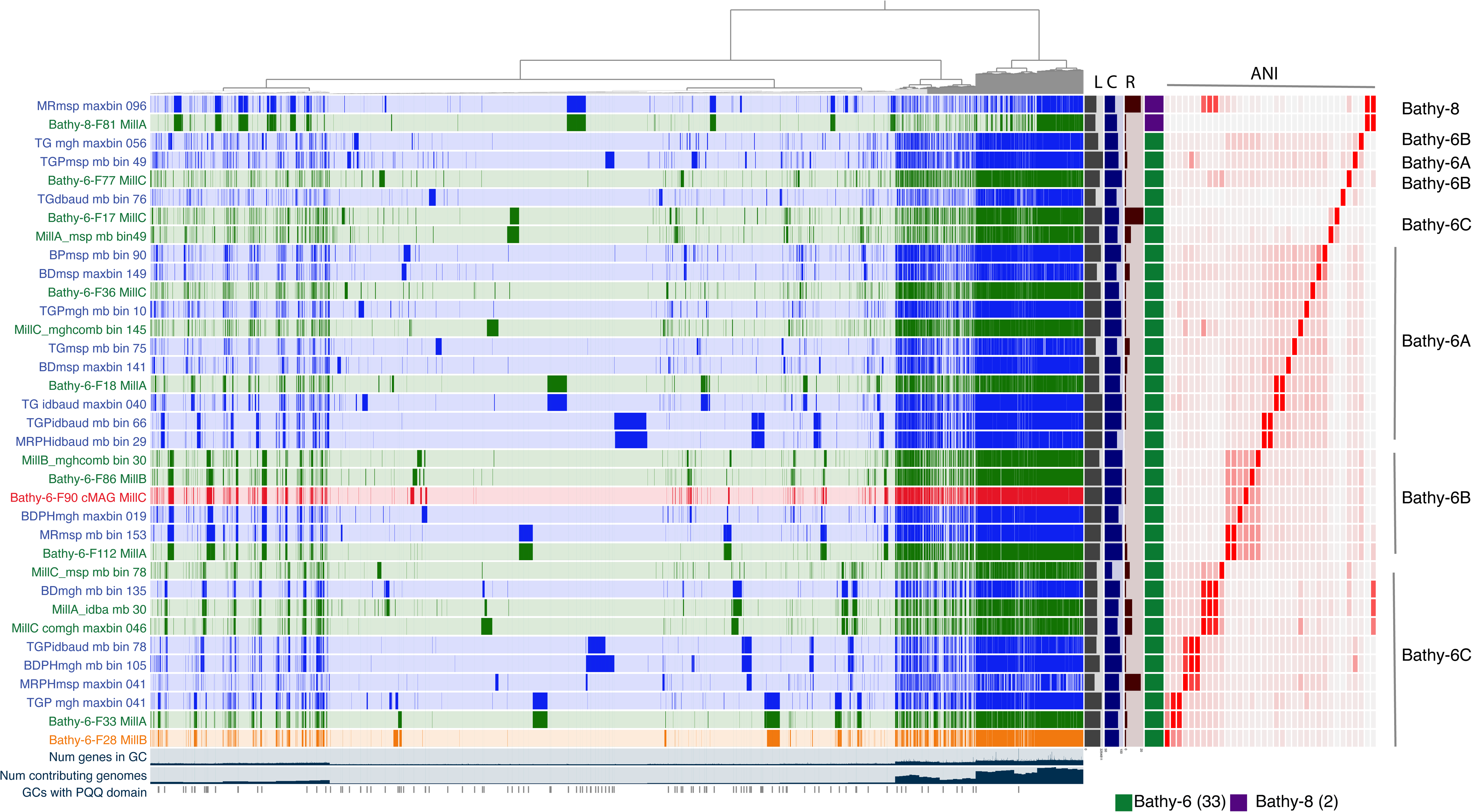
Pangenome generated in Anvi’o. Horizontal line 1 – 35 from the top represents a MAG and each vertical represents a gene cluster (GC). The MAGs are ordered by ANI identity. MAGs from the three Mills are shown in green except the cMAG which is highlighted in red and the high-quality MAG in one contig which is highlighted in orange. MAGs from the enrichment cultures are shown in blue. Line 36 and 37 horizontal lines display the number of genes in each GC and the number of genomes with genes in a GC. Line 38 shows the occurrence of PQQ-binding domains in the GCs. The dendrogram on the top represents the clustering of the GCs based on presence/absence. The bars on the right-side show C: % completion (50-100%), R: % redundancy (0-20%), L: total length of the genome and the Bathy 16S-rRNA lineage assigned in the phylogenetic tree in Figure 1. The ANI heatmap show ANI from 70% to 100%. The last column shows the Bathy clades from Fig. 1 and ANI similarity follows the clades in Fig. 1 except for MAGs found at the base of clades. Only MAGs generated in the current study are displayed. However, additional genomes from the genome taxonomy database (GTDB) were incorporated in the calculation of the pangenome and are included in Table S4 which contains the information on all gene clusters including annotation and sequences.

Manual refining of the MAGs resulted in a closed circular genome: Bathy6-F90_cMAG_MillC. The genome was 3,040,071 bp and contains 2928 CDS, 36 tRNA genes, two 5S rRNA genes, one 16S rRNA and one 23S rRNA gene (Table S1B and S3). All rRNA genes were found at separate loci. A second high quality MAG, Bathy6_F28_MillB was also in a single contig (Table1), but not closed. Of the remaining 33 MAGs, 6 were high quality, 23 medium quality and 4 low quality [29].

### Bathy-6 are anaerobic heterotrophic organisms

The cMAG was compared to the 34 MAGs reported here as well as 23 closely related MAGs from the GTDB (Fig. 2, Table S1, Table S4). This pangenome analysis identified 10,709 gene clusters (GCs) present in at least two genomes. Note that details for specific GC_IDs can be found in Tables S3 and S4. The core genome inferred from the pangenome analysis suggests the Bathyarchaeia assembled here are anerobic heterotrophic organisms and have a similar metabolic repertoire to Bathyarchaeia reported on earlier [2], including the recently described highly enriched cultures of Bathy-8 [8, 9]. A schematic of the predicted metabolism of Bathy6-F90_cMAG_MillC is shown in Fig. 3 and Fig. S2. The same overall metabolism was also inferred for the other Bathy-6 MAGs included in our analysis.

**Figure 3.**
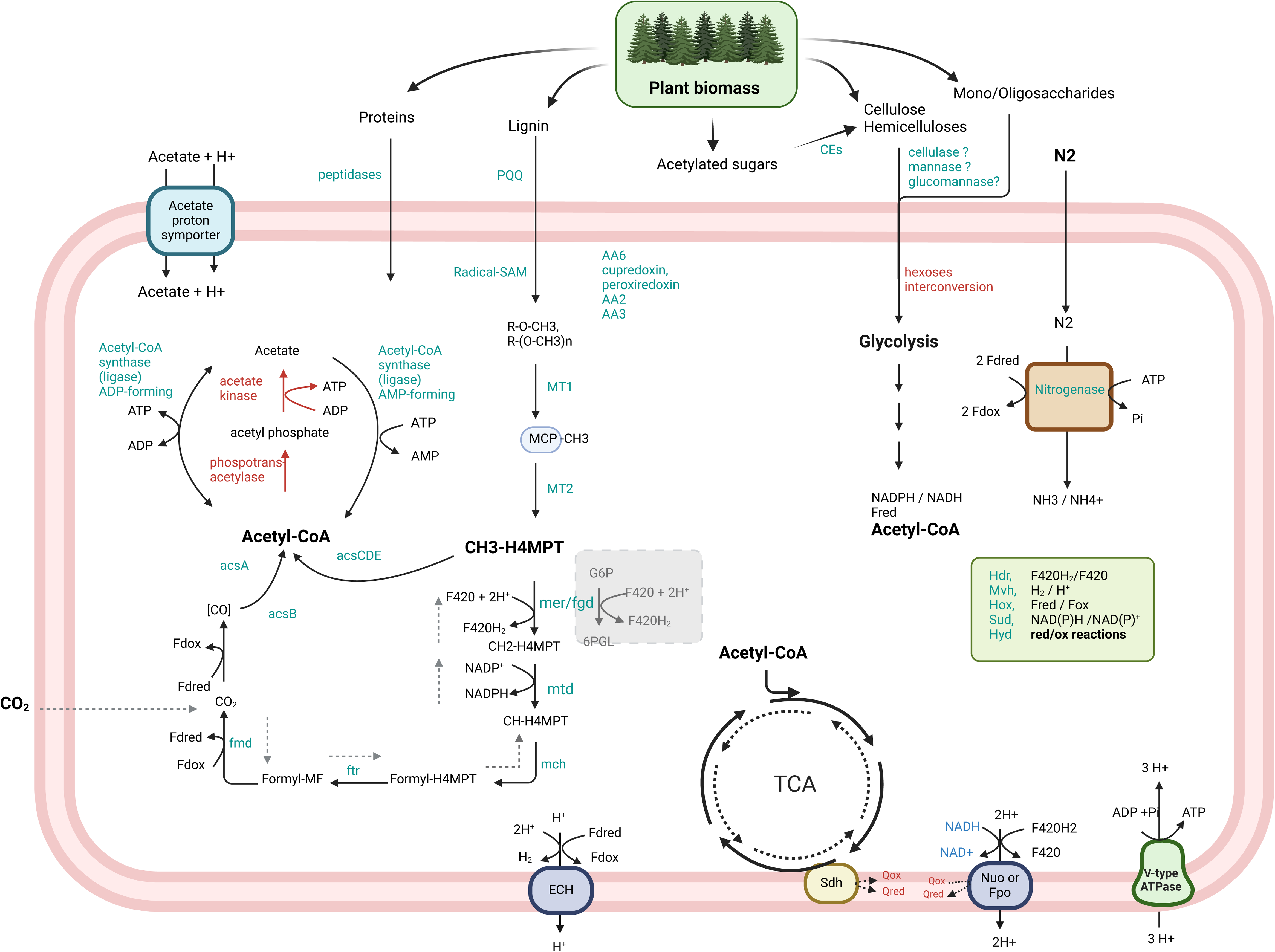
Overview of metabolic pathways reconstructed in the cMAG; Bathy6-F90_cMAG_MillC. Functions and proteins in green have been associated with a gene, while proteins in red have not been confidently assigned. Dotted lines indicate theoretically possible reactions, which have not been supported by the genome analysis. Protein abbreviations refer to the following gene ids in Table S1 with locus id from metaerg (GC ids in parenthesis): ECH: 551 - 559 (GC_00000178, GC_00000079, GC_00000535, GC_00001439, GC_00001128, GC_00001111, GC_00000834, GC_00000238, GC_00000742), FpoF: 2177 (GC_00000879), FpoJ/ NuoJ: 591 (GC_00001339), FpoBICDNHM/ NuoBICDNHM: 631 - 635 (GC_00002130, GC_00002053, GC_00002152, GC_00002167, GC_00002133), NuoGFE/ HoxUFE: 2708 - 2710 (GC_00001577, GC_00000659, GC_00001491), HoxHU/ MvhAG: 2706 - 2707 (GC_00001566, GC_00001601), MvhD/ hdrA: 2887 (GC_00000045), MvhB: 2886 (GC_00001129), HdrCB: 2744 - 2745 (GC_00000562, GC_00000381), H^+^ V-type ATP- synthase: 15 - 22, 24 (GC_00001573, GC_00001413, GC_00001548, GC_00001475, GC_00001334, GC_00001242, GC_00001374, GC_00001357), ADP-SCS/ ADP-ACS (Acetyl-CoA synthase (ligase) ADP-forming): 1955, 1102 (GC_00000166, GC_00000311), AMP-ACS (Acetyl-CoA synthase (ligase) AMP-forming): 423 (GC_00000360), ascABCDE: 2182 - 2186 (GC_00000082, GC_00000169, GC_00000340, GC_00000328, GC_00000188), MT1: 1643 (GC_00003051), MCP: 1919 (GC_00000134), MT2: 1920 (GC_00000430), Mer/Fgd : 448 (GC_00003096), Mtd/FolD: 2456 (GC_00000633), Mch: 954 (GC_00000326), Ftr: 2243, 1879 (GC_00000464, GC_00000374), FmdCFAC: 362 - 365 (GC_00000104, GC_00000038, GC_00000161, GC_00000744), FdhD: 366 (GC_00000729), SudAB: 671, 672 (GC_00000025,GC_00000891), SudD (paralog of A): 2736 (GC_00001894), Nitrogenase (NifHBDK): 2823 - 2826 (GC_00000249, GC_00001531, GC_00001597, GC_00001614), HydGBDA: 2711 – 2713(GC_00002346, GC_00002363, GC_00002610), predicted cellulase (endoglucanase, GH5_5): 835, 1425, 2894 (GC_00002139, GC_00003721), predicted mannase (GH5_7): 107, 152, 825 (GC_00003208, GC_00004040), deacetylases of unknown acetylated oligo/polysaccharides in the pulp material CE4: 2654 (not in pangenome) and CE14: 263, 1562 (GC_00001078, GC_00002987). For functions in the green box and a more detailed view of the energy metabolism see Supplementary Figure S3. The figure was drawn in Biorender.

All genes involved in glycolysis from glucose-1P or mannose-1P to pyruvate were identified in the cMAG and most of the Bathy-6 MAGs (Table S5) as described in detail in Supplementary text S5. The oxidative TCA-cycle (oTCA) has been reported incomplete (or missing) in many Bathyarchaeia MAGs [2, 9, 30, 31], but manual annotation of the proteins in the cMAG (Bathy6-F90_cMAG_MillC) allowed us to identify homologs of all TCA proteins (Fig. S3, Table S5). Using these genes as queries in blast searches identified many of the ‘missing’ genes also in the other MAGs, suggesting an active complete TCA pathway in Bathy- 6.

All the genomes had genes for the carbonyl- and methyl branch of the archaeal Wood Ljungdahl (WL) pathway. We did not detect homologs of methyl-coenzyme M reductase, ruling out methanogenic growth. The cMAG encoded all genes for a V-type ATP-synthase that utilizes the proton motive force (*pmf*) from the Fpo-like F420H2 dehydrogenase or Nuo-like NADH dehydrogenase, and energy-converting hydrogenase (ECH; Fig. 3, Fig. S2). The presence of the complete WL pathway is essential for, but not a conclusive marker of acetogenesis [32].

However, identification of the ECH and pyruvate synthase (POR), as well as ATP synthase, suggests acetogenesis from organic compounds is possible in Bathy-6.

Acetogenesis from H_2_ and external CO_2_ is not likely since we did not identify a bifurcating FeFe hydrogenase essential for providing low potential ferredoxins. For further discussion of these enzyme complexes and alternative roles, see Supplementary text S7.

### Bathy-6 archaea have genes for metabolising lignocellulose

The environmental distribution of Bathy-6 archaea suggest they are likely involved in digestion of some components of lignocellulose (plant biomass). In support of this we identified the genes recently shown to be involved in use of methyl- or methoxy groups common in lignin, producing methyltetrahydromethanoptein (CH3-H4MPT, Fig. 3) in Bathyarchaeia and other Archaea [2, 9, 33]. This two-step process involves transferring a methyl group from a methoxy compound via a methyl transferase (encoded by the MtgB gene) to a cobalamin-containing protein (MtgC), and then to tetrahydromethanopterin by a second methyltransferase (MtgA) [34]. The cMAG and all the Bathy-6 MAGs had the MtgA and MtgC genes (Fig. S4). Two to five homologs of the substrate specific *O*-demethylase (MtgB) identified in Bathy-8 [9] were also found in Bathy-6A and Bathy-6C MAGs (GC_00000053, GC_00000811, GC_00000883). Interestingly, the cMAG and the five other Bathy-6B MAGs lack MtgB homologs but carry a second MtgA homolog (GC_00003051), which we propose may perform substrate-specific demethylation. Several other genes were conserved in the MtgA-MtgC gene neighbourhood (Fig. S4). It is likely that some or all these enzymes participate in the methyl- or methoxy-group metabolism.

The methyltetrahydromethanopterin (CH3-H4MPT) produced by the methyltransferase system could be used by the CO dehydrogenase/acetyl-CoA synthase complex resulting in formation of acetyl-CoA, that could be converted to acetate with formation of ATP in the reversible reaction catalyzed by ADP-forming acetyl-CoA synthase [35] (Fig. 3). The cMAG and most MAGs also contained a gene for AMP-forming acetyl-CoA synthase (GC_00000360), but this enzyme is strongly directed towards acetate activation [35].

Additionally, the Bathy-6 MAGs had many genes involved in carbohydrate metabolism and the Bathy-6 cMAG had 125 genes with CAZyme annotations (Table S3). Among 25 glycosidases (GHs) encoded in the cMAG, six were assigned to the GH5-family (Fig. S5).

Pairwise-identity analyses and manual annotation suggested two GH5 subtypes (Table S3); three genes were most closely related to GH5_5 endoglucanases that may target cellulose and three genes were most closely related to GH5_7 mannanases [36]. All the genes were distantly related to characterized CAZymes and their function prediction should therefore be considered with caution. Also, the cMAG had one gene encoding an intracellular GH2 family enzyme and one extracellular GH113 enzyme (metaerg id 179; only observed in the cMAG), both predicted to be beta-D-mannosidases. Finally, four GH109 members might be involved in NAD+ dependent oxidation of various sugars and its derivatives. The other Bathy-6 MAGs encoded similar sets of GH-enzymes, while the Bathy-8 MAG only had one GH133 (Fig. S5). We identified 38 GC with pectin lyase domains (Table S3) which targets the pectin component of the plant cell wall [37].

These enzymes were predicted to be membrane associated with most having an extracellular pectin lyase domain. The cMAG has seven such genes from seven GCs.

The cMAG also had enzymes annotated as redox enzymes that act in conjunction with CAZymes or Auxiliary Activity (AA) oxidases [38] (GCs in Table S3); one AA2 catalase-peroxidase, one AA3 a long-chain alcohol peroxidase and 14 AA6 that may encode flavin-containing NADH-quinone oxidoreductases or flavodoxins (flavin-containing electron shuttles). Finally, many Bathyarchaeia have been proposed to use proteins and peptides as substrates (e.g. ref. [39]), and peptidases identified are reported in Supplementary text S3 and Fig. S5.

### Other genes involved in lignin metabolism

We identified 74 GCs in the pangenome predicted to be pyrroloquinoline quinone (PQQ)-dependent dehydrogenases (Fig 2, Table S4). In *Methanosarcina* PQQ-binding proteins have been shown to be involved in extracellular electron transport in humic acid-dependent respiratory growth [40] and we therefore hypothesize that these enzymes in Bathyarchaeia interact in a similar way with lignin. Interestingly, different PQQ-GCs were abundant in the three Bathy-6 lineages depicted in Fig. 1. PQQ-binding proteins from GC_00000001 were abundant in the Bathy-6 clade A where the genomes had on average 24 copies of this gene (Fig. S6).

GC_00000010 and GC_00000981 contained PQQ-dehydrogenase-genes abundant in clade B (including the cMAG) with average 11 and 4 copies, respectively. In clade C GC_00000116, GC_00001367, and GC_00001610 are abundant. The cMAG had 46 PQQ-domain-encoding genes and of these 24 were from GC_00000010 (Fig. S7). Almost all the predicted PQQ-dehydrogenases had a Sec dependent signal peptide and thus are predicted to be extracellular. We also identified a possible PQQ-synthase gene (GC_00000466) in the cMAG and in most of the Bathy-6 MAGs (Table S4). This gene has a PqqA peptide cyclase domain and a structure most similar to ‘coenzyme PQQ synthesis protein E’ from *Methylobacterium extorquens* (PDBe 6c8v; 27% identity and 100% confidence by Phyre2) [41].

Another abundant annotation in the pangenome was ‘Radical SAM domain’-proteins (Table S4). Radical SAM enzymes catalyze a wide variety of reactions such as methylations, isomerization, and reduction via protein radical formation [42], and are likely candidates to be involved in metabolism of lignin. Twenty-five GCs had the annotation COG1032 (Radical SAM superfamily enzyme YgiQ) and were assigned to Megacluster-2 in the Radical SAM database, which contains vitamin B12-dependent enzymes [43]. Each MAG contained 7-20 of these genes distributed in 4 - 14 GCs (Table 4). The cMAG had 11 such genes from 10 GCs (Fig. S7). In addition, 38 Radical SAM-GCs were assigned to Megacluster-1-1 in the Radical SAM database and had the COG annotation COG0535 (Radical SAM superfamily maturase) (Table S4). The cMAG had 17 such genes from 17 GCs. Several genes involved in B12-synthesis (see below) and the PQQ-synthase mentioned above were found in this category.

### Bathy-6 have a positive effect on the community through N-fixation and vitamin B12 production

The cMAG and several of the Bathy-6 MAGs had genes for a Mo-nitrogenase (Fig. 3, Fig. S8). The genomic region contained the *nifHBDK* genes, and the conserved active site residues in *nifD* [44] were also conserved in the Bathyarchaeia homologs. Homologs of *nifE* and *nifN* were not found in the cMAG (or any other MAG). This has also been observed in early branching nitrogenase lineages [44]. Phylogenetic analysis of the *nifD* and *nifK* proteins revealed that the Bathyarchaeia nitrogenase clustered with the deeply branching *Chloroflexaceae* (Clfx clade in [44]) (Fig. S9). Moreover, the Bathyarchaeia nitrogenase and the Clfx have the gene arrangement *nifHBDK* instead of the *nifHDK* observed in other lineages.

Most of the genes for synthesis of vitamin B12 from glutamate could be identified in the Bathy-6 MAGs, including the cMAG where they were found at three loci (Fig. S10). We also detected homologs of some of the genes involved in synthesis of the lower ligand [45]; BzaCDE. One of these genes was at a B12-synthesis genomic locus in the cMAG (gene id 2298). The Bathy-6 genomes also encoded a putative BtuFCD ABC transporter, importing cobinamides from the environment. An adenosylcobinamide hydrolase (CbiZ) was found next to the transporter, supporting its role in cobinamide salvage. Most of the Bathy-8 MAGs included in the pangenome analysis only have genes for the BtuFCD transporter and homologs of the enzymes in the salvage pathway (Fig. S10). The Bathy-8 genomes from the two highly enriched cultures, however, carry all of the genes identified in the cMAG and the Bathy-6 MAGs [8, 9]. The missing genes in the other Bathy-8 genomes could therefore be due to the MAGs being incomplete. Additional high-quality genomes from the Bathy-8 lineage are needed to confirm this.

### Lateral gene transfer and genomic islands

The numerous PQQ-dehydrogenase genes in the Bathy-6 genomes were likely acquired through lateral gene transfer (LGT) since these proteins have a scattered distribution in Archaea [40]. We were unable to predict the exact donor lineage of these genes since the Bathyarchaeia proteins are highly divergent and BLAST searches only detected significant matches of within Bathyarchaeia. Among Archaea in the IMG database [46], most PQQ-domains (pfam13360) are found in Halobacteria, Methanosarcina, and Bathyarchaeia. The cMAG and our MAG in one contig (Bathy6-F28_MillB), are the two archaeal genomes in IMG with the highest number of pfam13360-protein encoding genes.

The phylogenetic analysis also suggested that nitrogenase genes discussed above were acquired by the ancestor of Bathy-6 from an unknown lineage (Fig. S9). More recent LGT events were also detected. Phylogenetic analysis of 135 genes from the cMAG with no match in the other genomes included in the pangenome analysis, suggested that 80 have been acquired by LGT (Table S3). Some of these genes were found co-localized and a comparison of the high-quality MAGs (Table 1) to the cMAG revealed four large genomic regions (GRs) with few or no matches to other Bathy-6 (Fig. S7, Table S3). Similar island regions were observed in the Bathy6_F28_MillB MAG, the second MAG in a single contig, suggesting this is a common feature of these genomes (Fig. S11).

Genomic region 1 (GR-1) was ∼99kb with 95 CDSs, and contained a 46,769 bp provirus identified by Virsorter [47] with typical virus genes such as a terminase, integrase, and phage structural genes (Fig. S12). Virfam [48] suggested the virus belongs to the *Siphoviridae* based on its head-neck-tail proteins. Phylogenetic analyses showed that the virus region contained genes widespread in Bathyarchaeia interspersed with genes from distant lineages (Table S3), indicating that this virus is common in Bathyarchaeia. The other genomic regions are described in Supplementary text S4.

## DISCUSSION

### Long read sequencing and ‘low’ diversity produced high quality MAGs

Most Bathyarchaeia lineages are only known as MAGs, often of low quality and fragmented into numerous contigs (average 200 contigs for MAGs classified as Bathyarchaeia in Genbank, February 2024). Recently a 2.15 Mb cMAG was recovered from a highly enriched Bathy-8 culture [8]. Here we assembled eleven high quality MAGs, including one closed 2.8 Mb chromosome (Bathy6-F90_cMAG_MillC) and a genome represented by a single 2.7 Mb contig (Bathy6-F28_MillB). This was enabled by using long-read sequencing and sequencing metagenomes with relatively high abundance of a few Bathyarchaeia genomes (Table 1, Supplementary text S6). The larger size of the Bathy-6 cMAG and MAGs compared to the Bathy-8 cMAG, indicate that Bathy-6 are more metabolically versatile than Bathy-8.

### Lateral gene transfer and gene expansions shape the Bathy-6 genomes

We found that both LGT (with both bacteria and archaea; Table S3) and gene family expansions through duplication have been particularly important during the evolution of these organisms.

These evolutionary mechanisms have also been highlighted in other archaea from Thermoproteota [49]. The numerous PQQ-dependent dehydrogenases illustrate the importance of both evolutionary processes. The patchy phylogenetic distribution of these genes in Archaea suggest that they were acquired by LGT, while the distribution within each Bathy-6 genome with multiple copies of genes from the same GC suggests recent independent expansions within each lineage, likely due to within genome duplication.

The flexible genome was also evident when we investigated the proteins involved in core metabolic pathways. For instance, while most of the MAGs had homologs of the TCA enzyme NADPH isocitrate/isopropylmalate dehydrogenase (EC:1.1.1.41/EC:1.1.1.85, COG0473; GC_00000347), two MAGs did not have this gene and instead encoded another NADPH isocitrate dehydrogenase (EC:1.1.1.42, COG0538; GC_00005384) (Table S5, and Supplementary text S7 for additional examples.)

### Bathy-6 is common where lignin is present and contributes important community functions

The Bathy-6 lineage was the most abundant Bathyarchaeia lineage in the pulp mills and the poplar hydrolysate enrichment cultures, with a total of 33 Bathy-6 MAGs compared to only two Bathy-8 genomes. This suggests that the Bathy-6 lineage is better adapted to life in these anaerobic lignocellulose rich environments. Several other observations also support the importance of the Bathy-6 lineage in degradation of plant material. Bathy-6 are common in terrestrial anaerobic environments with decaying plant material such as the permafrost active layer and termite guts (Fig. 1). Furthermore, in Mill C, the abundance of Bathy-6 was positively correlated with color concentration in the final effluent, where the color was mainly caused by lignin derived compounds [13]. Most of the MAGs obtained here contain the *O*-demethylase system recently proposed by ref. [9] which is a possible reason for their success in communities enriched in lignin compounds. However, the *O*-demethylase system was also present in Bathy-8 organisms; therefore, these genes alone cannot explain the success of Bathy-6.

One set of enzymes that may contribute to the success of Bathy-6 in these environments compared to Bathy-8 are CAZyme-encoding genes. In particular, the Bathy-6 MAGs encode several CAZymes targeting cellulose and hemicelluloses (Fig. S5) not found in Bathy-8, including the Bathy-8 recently linked to lignin degradation [8, 9]. The Bathy-6 metabolism is therefore closely linked not only to lignin degradation but also to degradation of polysaccharides in the pulp and paper mill wastes. The fact that Bathyarchaeia were less abundant in the cellulose-enriched cultures (Table1), suggests that the enzymatic set carried by the cMAG and other Bathy-6 MAGs have a preference for hemicelluloses and mannans in particular.

Endoglucanases encoded in the cMAG may act on glucose-containing heteropolysaccharides, synergistically enhance the rates of action of other glycosidases [50], and/or possibly help improve access to lignin. Taken together, this shows that the Bathy-6 metabolism is well suited for life in a pulp and paper mill bioreactor, a beaver’s intestines, or a moose rumen.

The Bathy-6 archaea may also be more self-sufficient than the Bathy-8 organisms. We found that the cMAG and most of the other Bathy-6 MAGs encode an almost complete anaerobic pathway for cobalamin or vitamin B12 synthesis (Fig. S8), which is an important co-factor of many enzymes including those involved in the *O*-demethylase system and the W-L pathway. This pathway is missing in most Bathy-8 MAGs. Vitamin B12 is synthesized by a restricted number of prokaryotic organisms and mostly Bacteria [51]. Among archaea, only Thaumarchaeota have been reported to have the enzymatic machinery for B12 synthesis [52].

Only two of 23 B12 synthesis genes were not identified in the cMAG, and Hou *et al*. [2] also identified many of these genes in other Bathy-6 MAGs. The ‘missing steps’ in the pathway are likely performed by yet unidentified enzymes since several missing proteins were also observed in the B12-synthesis pathway of *Thermosipho* spp., which are known to produce vitamin B12 [53].

It is also likely that many Bathy-6 organisms can reduce N_2_ to bioavailable ammonium. Recently, the presence of a nitrogenase gene in another Bathyarchaeia MAG was reported [31], however, closer inspection revealed that these were short contigs (WUQU01000012; 1725 bp, WUQU01000437; 2043 bp) with 97% -100% identity to proteins from *Caldicellulosiruptor* spp, and were likely contaminations. The nitrogenase genes identified here are therefore, to our knowledge, the first confident identification of a full nitrogenase gene cluster in Bathyarchaeia. Moreover, it is the first nitrogenase gene cluster found in an archaeon outside of a methanogen lineage.

The Bathy-6 lineage will therefore likely positively affect their microbial communities by contributing vitamin B12 and ammonium. In agreement with this, correlation analyses of 16S rRNA ASV data from the Mills identified several Bathy-6 ASVs in community modules operating under different environmental conditions [13] (Fig. S1). For instance, in Mill A 16S rRNA ASVs representing Bathy8-F81_MillA and Bathy6-F33_MillA were linked to stable conditions, while ASVs representing Bathy6-F28_MillA, Bathy6-F86_MillA and Bathy6-F112_MillA were linked to stressful conditions [13]. In Mill B both MAGs that could be linked to 16S data (Bathy6-F28_MillB and Bathy6-F86_MillB) were correlated to high COD removal efficiency, while in Mill C 16S rRNA from the Bathyarchaeia MAGs (including the cMAG) were linked to color formation as discussed above. Genes in the large accessory genome (Figure 2) are likely responsible for different Bathyarchaeia populations thriving under different conditions.

### A possible mechanism for lignin modification or degradation by Bathy-6

The high abundance of PQQ-dehydrogenase-like genes in the MAGs suggest their metabolic importance. In *Methanosarcina,* PQQ-binding proteins were recently shown to be involved in extracellular electron transport in humic acid-dependent respiratory growth [40]. In Bathyarchaeia, we propose that these proteins are involved in redox reactions with lignin and electron transfer. The reduction potential of PQQ is +90 mV [36], making them strong electron acceptors capable of electron acquisition from various donors - for example, different lignin moieties. In the cMAG, several of the PQQ proteins are multidomain proteins bearing more than one PQQ domains as well as, in many cases, immunoglobulin-like domains possibly involved in substrate binding. We did not find any known quinone synthesis pathways in Bathyarchaeia, suggesting that their quinones are either distinct or, perhaps more likely, that they are absent, as seen in acetogens and some methanogens [54]. Moreover, PQQ and lignin are both structurally similar to quinones and could act as intermediate acceptor of electrons provided by the TCA and the Fpo/Nuo-like complexes.

The AA2, AA3, and AA6 oxidases identified are also likely are involved in metabolism of lignocellulose. Characterized AA2 (catalase-peroxidase) and AA3 (long-chain-alcohol oxidase) use oxygen or reactive oxygen species (ROS). Since Bathy-6 inhabit anoxic environments, it is likely that some of these enzymes interact with other compounds such as lignin or lignin degradation products. AA6 are flavin-containing proteins involved in electron shuttling (flavodoxins) or reduction of various electron acceptors using NADH as the electron donor.

Three small intracellular iron-sulfur proteins (GC_00000897, GC_00001462, GC_0000199; Table S3), that have Rieske domains most closely related to the Rieske domains of various alkene monooxygenases [55] are also likely involved in electron transfer. Importantly, the oxygenase subunits of these alkene monooxygenases were not identified, thus the compound accepting electrons from Rieske proteins remains undefined. In the cMAG, one Rieske protein-encoding gene is co-located with two genes encoding AA6 homologs (metaerg ids 685, 686, and 678 in Table S3), suggesting possible electron shuttling from AA6 to an unknown electron acceptor through the Rieske protein. Other possible electron donors to Rieske are NADH-cytochrome b5 reductase (metaerg id 426), NADH:flavin oxidoreductase, and cytochrome b5 (metaerg id 88 - 89), all of which might acquire electrons from unknown quinones or PQQ-domain proteins.

We were not able to confidently identify the Bathy-6 cells terminal electron acceptor, nor the enzymatic machinery catalyzing terminal electron. One possibility is that internal proteins act as the electron sink. Radical-SAM proteins, which are abundant in the Bathyarchaeia genomes, could act as such acceptors since their ferredoxin domain must be reduced to form active 5′-deoxyadenosyl radicals. Enzymes using these radicals might be involved in lignin decomposition by making it more accessible to oxidation by PQQ, AA2, or AA3 as well as to demethylation and the formation of methoxy-compounds – the substrates for the methyltransferases MT1 and MT2.

It is also possible that Bathy-6 acquire traces of oxygen from the environment or during detoxification of ROS. Low amounts of oxygen may not inhibit these anaerobic microorganisms but be enough for lignin degradation. This oxygen could be bound and transferred to oxygen-requiring enzymes by bacteriohemerythrin (encoded by GC_00003974).

### Concluding remarks

The analysis of the complete Bathyarchaeia genome along with several high-quality MAGs allowed us to make predictions of their metabolism with more certainty. We identified all the required components for heterotrophic growth on sugars and lignin. The large genome size, presence of a full vitamin B12 synthesis pathway and a nitrogenase gene suggest that the Bathy-6 organisms studied here are largely self-sufficient and would explain their role as keystone species in the community. Careful manual annotation of genes involved in energy metabolism suggest acetogenesis from organic compounds is possible, however, not from H_2_ and external CO_2._ The nature of the environments where these Bathy-6 thrive, as well as their cohabitation with methanogens also make the possibility of lithoautotrophic acetogenesis unlikely. We suggest that their gene repertoire rather reflects a nitrogen-low diet and a metabolism heavily dependent on enzymes that use vitamin B12 dependent radical SAM enzymes. The latter fits well with our hypothesis that lignin and carbohydrates are their main source of energy and carbon.

## Data availability

Accession numbers for reads and assemblies are listed in Table S1. Fasta files of all the MAGs and phylogenetic trees of genes in the GRs are available on FigShare 10.6084/m9.figshare.26304676 and 10.6084/m9.figshare.26307589. High-quality MAGs have been submitted to Genbank under bioproject PRJNA916529. Two MAGs, the closed MAG (Bathy6-F90_cMAG_MillC and a MAG in one contig, Bathy6_F28_MillB, were submitted to and annotated in the IMG database [46]; taxon id 8000479782 and 8000476869, respectively.

## Acknowledgement

The project was funded by the Genome Canada Synbiomics project (#10405) with support from Ontario Genomics, Genome Quebec, and Genome British Columbia. Additional funding was provided by NSERC through a Discovery Grant to EAE. We are indebted to the personnel of the three mills for their help with sampling. We want to thank Mike Lacourt, Mabel Wong and Shen Guo for establishing and/or maintaining enrichment cultures. We’d like to thank Emma Master for support and guidance. Dr. Julien Lossouarn for help with identifying the provirus sequence.

