## Supplementary text for "High quality Bathyarchaeia MAGs from lignocellulose-impacted environments elucidate metabolism and evolutionary mechanisms"

### **Supplementary Text S1: DNA extraction, sequencing, metagenome assembly and binning.**

Sample collection and DNA extraction was performed as described in Meyer et al. [1]. Shotgun metagenomic libraries with an insert size of 500bp were constructed from six samples (Table S1A) and sequenced on an Illumina NextSeq 500 sequencer generating paired-end read of 150nt at the Centre for the Analysis of Genome Evolution and Function (CAGEF) at the University of Toronto, Canada. One sample from each mill was also sequenced using PacBio CS at Genome Quebec, Montreal, Canada (Table S1A). The Illumina reads were quality trimmed and assembled using the Anvi’o v. 6.2 metagenomic snakemake workflow [2]. This pipeline does read quality control using the methods developed in Minoche et al. [3]. The quality trimmed reads were assembled using three different assemblers: SPAdes v. 3.12.0 [4], MEGAHITv. 1.2.9 [5] and IDBA-UD v.1.1.3 [6]. The assemblies were performed on the samples separately and combined in co-assemblies. Finally, PacBio reads were used in hybrid assemblies with the Illumina reads from the same sample using SPAdes v3.14.1 (in metaspades mode). The metagenomic sequencing of the enrichment cultures are described in references [7] and [8]. The reads from these samples were also assembled using SPAdes v. 3.12.0, IDBA-UD v.1.1.3 and MEGAHIT v. 1.2.9. Binning of all the assemblies was performed using metaBAT 2 v2.15 [9] and MaxBin2 v.2.2.7 [10], and bins from the same mill or culture were dereplicated using dRep v.2.5.4. [11].

### **Supplementary Text S2: Refinement and annotation of MAGs**

MAGs with a 16S rRNA sequence matching an ASV from the amplicon datasets were renamed using the Bathyarchaeia lineage, ASV-id, and the mill they originated from (e.g. Bathy6-F18_MillA). MAGs from the mills with few contigs (< 30) were selected for further refinement using *de novo* assembly of the contigs in Geneious Prime ([www.geneious.com](https://www.geneious.com)), SSPACE (Boetzer *et al.*, 2011), the ra2.py script (<https://ggkbase-help.berkeley.edu/genome_curation/genome-curation/>), as well as mapping of PacBio CCS reads using Minimap2 in Geneious Prime and mapping of Illumina reads using the Geneious mapper (with the “only paired reads that map nearby” option). One MAG, originally in four contigs from the hybrid metaspades assembly of MillC, was closed by the assembly of contigs in Geneious Prime (cMAG). The contig junctions of the cMAG were confirmed by mapping pacbio CCS reads and contigs from a Flye v. 2.9.2 assembly (of all MillC pacbio reads) using Minimap2 in Geneious Prime. Replication origins of the cMAG and one MAG in a single contig was predicted by Ori-Finder 2022 [13]. For all MAGs, possible contaminating contigs that had no coding sequence (CDS) assigned to Bathyarchaeia by the metaerg annotation pipeline [14] and that also did not have homologs in other Bathyarchaeia MAGs were removed from the MAGs.

In addition to annotation using metaerg [14], the cMAG was annotated using dbCAN3 [15], TMHMM 2.0 [16], SignalP 5.0 [17], Phyre2 [18], MEROPS [19] and Interproscan [20]. Proteins in suspected provirus regions were compared to the NCBI virus database using BlastP. Provirus classification was performed using Virfam [21].

### **Supplementary Text S3: Additional results from characterization of the cMAG genome**

The cMAG carried 257 genes with no close homologs in the other MAGs in the pangenome analysis (Supplementary Table S3). Hypothetical proteins were the most numerous (N=163), while the second most common annotation was glycosyl transferase (N=13), mainly from the GT4 and GT2 families.

Among CAZymes not discussed in the main text, two carbohydrate esterases (CEs) in the cMAG belonged to CE14 family and one to CE4 family (Supplementary Table S3). All characterized archaeal CE14 are involved in N-acetylglucosamine deacetylation. For the cMAG we propose a similar function of these enzymes. No polysaccharide lyases were found.

We identified 25 peptidases in the cMAG by searching the MEROPS database, 8 of these were predicted to be extracellular (Supplementary Figure S5) and might be involved in degradation of proteins in the environment. The most abundant family observed was the S1 peptidase family, which was found in all the MAGs (Supplementary Figure S5). The cMAG also encoded peptidases from the S08 family, the PepG1 family and carboxypeptidases. Moreover, we identified possible peptide transporters in all the genomes (Supplementary Table S4).

### **Supplementary Text S4: Genomic regions 2, 3 and 4 in the cMAG**

Genomic region 2 (GR-2) was composed of 40 genes and contained (remnants of) a mobile genetic element with genes encoding proteins involved in DNA metabolism including a reverse transcriptase-like protein and a CRISPR-Cas protein. Genomic region 3 (GR-3) was less well defined but contained at least 14 genes, including three that encode GT family 4 proteins and a CE family 4 protein. Genomic regions 4 and 5 (GR4, GR5) were also enriched in CAZy-genes (e.g. GTs) and other genes annotated as transferases (acetyl- and methyl-transferases) with diverse taxonomic distribution, suggesting that these genes have been frequently lost and acquired by these organisms.

### **Supplementary Text S5: Identification of genes involved in glycolysis.**

All genes involved in glycolysis from glucose-1P or mannose-1P to pyruvate were identified in the cMAG and most of the Bathy-6 MAGs (Supplementary Table S5). A specific glucokinase converting glucose to glucose-1P was only observed in Bathy-8 MAGs (Gene cluster ID [GC]: GC_00003139). Note that details for specific GCs can be found in Tables S3 and S4). Archaea commonly use other kinases for this step, often with a broader substrate range [22, 23], and this is likely also the case for Bathy-6. A candidate gene for this function was the sugar or nucleoside kinase (COG0524) encoded by the genes in GC_00000937 (Supplementary Table S5), exclusively found in MAGs that do not carry the GC_00003139 glucokinase. In support of this, homologous haloarchaeal kinase was predicted to be a fructokinase based on genomic and proteomic data [22]. A phosphoglucose/phosphomannose isomerase (GC_00000674) connects mannose oxidation and the glycolysis. The cMAG and most of the MAGs had a pyruvate synthase (POR; Supplementary Table S5) which catalyzes the ferredoxin reducing decarboxylation of pyruvate to acetyl-CoA – a key reaction for production of low-potential reduced ferredoxins.

### **Supplementary Text S6: Genome size and structure in Bathy-6**

The cMAG (2.8 Mb) and high-quality MAGs (2.5 – 3.0Mb) are considerably larger than most Bathyarchaeia genomes in Genbank (average 1.6Mb in January 2023). One other cMAG is available from a highly enriched Bathy-8 organism which is 2.15 Mb [24]. This is also smaller than the Bathy6-F90_cMAG_MillC and Bathy6-F28MillB, which suggests that Bathy-6 genomes are larger than those in Bathy-8, and probably also other Bathyarchaeia.

The genome structures we observed give some hints to why they appear to be particularly difficult to assemble from metagenomic data. Both Bathy6-F90_cMAG_MillC and Bathy6-F28_MillB contain several regions with few or no matches in other Bathyarchaeia, suggesting these are region of high diversity between closely related genomes that can complicate assembly. All the MAGs assembled by us also have large gene families with many copies of very similar genes (the PQQ-domain proteins and the vitamin B12 dependent Radical SAM enzymes) which also may interfere with assembly.

### **Supplementary Text S7: Shared phylotypes across lignin impacted sites.**

The TG culture originated from an inoculum from MillA in 2010. Two MAGs from the TG culture were very similar (>0.98 ANI) to MAGs from MillA (Figure 1 and 2, Supplementary Table S2); Bathy6-F33_MillA and TGP_mgh_maxbin.041 have ANI of 0.98 and Bathy6-F18_MillA and TG_idbaud_maxbin.040 an ANI at 0.99. However, we also found highly similar lineages between MillA and MAGs from the moose rumen culture (MR) and beaver droppings (BD), and between the three cultures (Supplementary Table S2). Among the mills, MillA and MillC shared one lineage, with MillA_idba_mb_bin.30 and MillC_comgh_maxbin.046 having an ANI of 0.96. Interestingly, a similar genome was also observed in the beaver droppings enrichment; BDmgh_mb_bin_135 and MillA_idba_mb_bin.30 have an ANI at 0.99. Taken together, this demonstrates that the mills, beavers, and moose contain Bathyarchaeia that are very similar. This may mean that Bathyarchaeia have recently (in evolutionary terms) expanded into these environments.

### **Supplementary Text S7 Carbon and energy metabolism in Bathy-6**

*A complete TCA cycle with a diverse set of enzymes.* We found that the MAGs have different sets of glycolysis- and TCA cycle enzymes (Supplementary Figure S4 and Table S5 for GC-ids mentioned below). For instance, while most of the MAGs had homologs of an NADPH:isocitrate/isopropylmalate dehydrogenase (EC:1.1.1.42/EC:1.1.1.85, COG0473; GC_00000347), two MAGs are missing this gene and instead encode another NADPH isocitrate dehydrogenase (EC:1.1.1.42, COG0538; GC_00005384). Six of the MAGs from the 6B and 6C clade had a ‘classical’ *sucD* and *sucC* combination (EC:6.2.1.5; GC_00002664 and GC_00002457), while these genes were not found in the cMAG and most of the other MAGs. Instead, we identified two homologs of other succinyl-CoA synthases in the cMAG. The first had a domain structures similar to the two subunit *Thermococcus kodakaraensis* [23, 25] succinyl-CoA synthase (SCS) (GC_00000166, GC_00000311). The second resemble the multidomain acetyl-CoA synthase (ACS) (GC_00000078) from the amitochondriate eukaryote *Giardia lamblia* (Sánchez et al., 2000). Similarly, 12 of the MAGs from 6B and 6C have a two-subunit class II fumarate hydratase (EC:4.2.1.2; GC_00001815, GC_00001804) while the cMAG and the other MAGs have a fused protein (EC:4.3.1.1; GC_00001135). In support of the Bathy6-F90_cMAG_MillC protein being a fumarate hydratase, its gene was found next to the succinate dehydrogenase genes (GC_00001921, GC_00001923), which catalyzes the preceding reaction in the oTCA. Taken together this suggests that the oTCA is present in all Bathy-6 and shows a high degree of plasticity with its enzymes frequently being substituted with homologous or analogous proteins. Finally, we cannot discount the possibility of reversed operation of TCA i.e. roTCA [26]. If this is true, the Bathy-6 lineage might be also capable for chemoorganoautotrophy as it was recently proposed for ANME-2 and Bathy-8 representatives [27].

*Carbon and energy metabolism in Bathy-6.* The energy conservation in the cMAG and the other Bathy-6 MAGs, may be achieved by acetogenesis from methyl- or methoxy compounds from lignin, glycolysis and TCA (all are substrate-level phosphorylation) and by V-type ATP-synthase (Figure 2). Among three possible complexes that may generate *pmf* for ATP-synthetase - F420H_2_ dehydrogenase (Fpo), NADH:ubiquinone oxidoreductase (Nuo) and energy converting hydrogenase (ECH) - Fpo and ECH are the most likely.

Fpo and Nuo are homologous protein complexes capable of exporting protons while reducing membrane electron carriers by oxidation of F420H2 or NADH, respectively [28]. Theses complexes have common electron transferring and membrane subunits and can be distinguished by the subunits involved in electron donor reduction. Since both F420H_2_ dehydrogenase and NADH dehydrogenase were identified in the cMAG and the genes encoding catalytic subunits of both of them are distantly located in the genome from their respective membrane-subunits-genes, it is impossible to determine which complex is active (Supplementary Figure S3). The current consensus is that Nuo is absent in Archaea [29] making the Fpo a more likely option. If this is indeed the case, the NADH dehydrogenase subunits (NuoGFE) could be a part of a cytoplasmic hydrogenase Hox (HoxUFE in this case). This is supported by the gene context analysis, since the genes encoding other putative Hox subunits (HY, metaerg locus id 2706-2707 in Table S3) are co-located next to the *HoxUFE* genes (metaerg locus id 2708-2710). Finally, the NuoGFE/HoxUFE subunits are also homologous to subunits of the bifurcating hydrogenase used by acetogens for producing low-potential ferredoxins [30]; however, the catalytic subunit (metaerg id 2708, Supplementary Table S3) is shorter and lacks a FeFe hydrogenase domain needed for it to function as a bifurcating hydrogenase and thus making it impossible for the Bathy-6 organism associated with the cMAG to grow by acetogenesis from external CO_2_. ECH is an energy-converting hydrogenase exporting a proton while reducing two protons with electrons from reduced ferredoxin which could be involved in *pmf* generation in various metabolisms. Since no quinone biosynthesis genes were found, it is possible that Fpo-like or Nuo-like complexes in Bathy-6 use electron acceptors other than quinones, while keeping their function in exporting protons i.e. in *pmf* generation. The cMAG also encoded homologs of a cytoplasmic hydrogenase (Hyd) as well as cytoplasmic subunits of heterodisulfide reductase Hdr (HdrABC) and sulfide dehydrogenase Sud (Supplementary Figure S3). We suggest they may replenish reducing equivalents and/or be responsible for production of H_2_ or reduced sulfur compounds. The Hyd[GB]DA genes (metaerg ids 2711-2713) are located next to (but on the opposite strand) the predicted *HoxHYUFE* assuming their possible co-action.

### **Supplementary Figures**

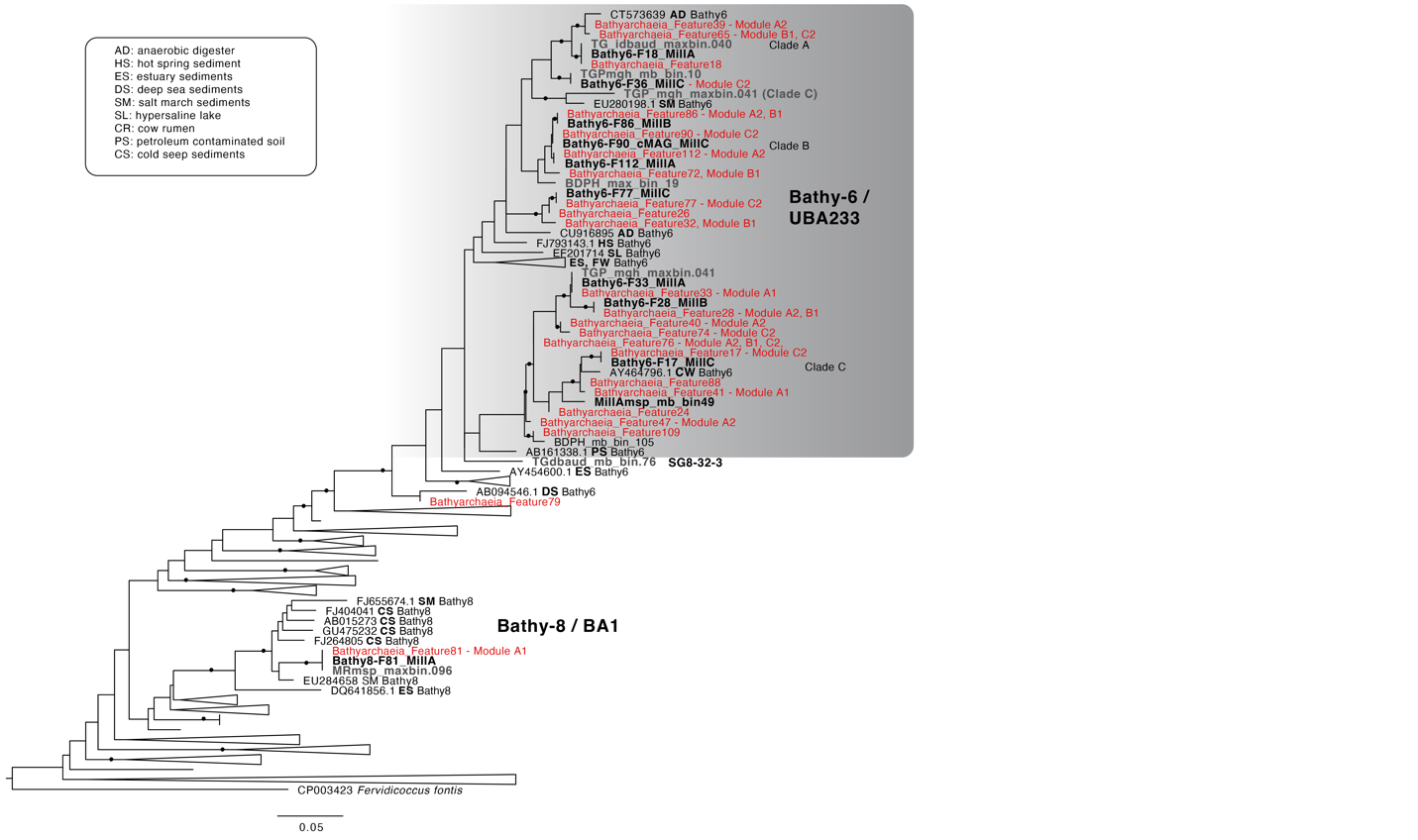

**Figure S1.** **Maximum likelihood phylogenetic tree of 16S rRNA genes from the Bathyarchaeia MAGs and 16S rRNA amplicon sequence variants (ASVs) labelled Bathyarchaeia_Fearture_xxx from [1].** If the ASV was included in the functional modules identified in [1] this is also indicated. The A1 and B1 module was correlated with stable conditions while module A2. In addition, Bathyarchaeia ASVs in module C2 were correlated with color in the effluent. Representative 16S rRNA genes from Bathyarchaeia lineages, MCGs, were obtained from [31]. The sequences were aligned in Geneious Prime using MAFFT [32]. The Maximum likelihood phylogeny was obtained using RAxML [33] in Geneious Prime with the GTR + GAMMA model and 100 bootstrap replicates. Black circles branches indicate bootstrap support values >= 70%.

 
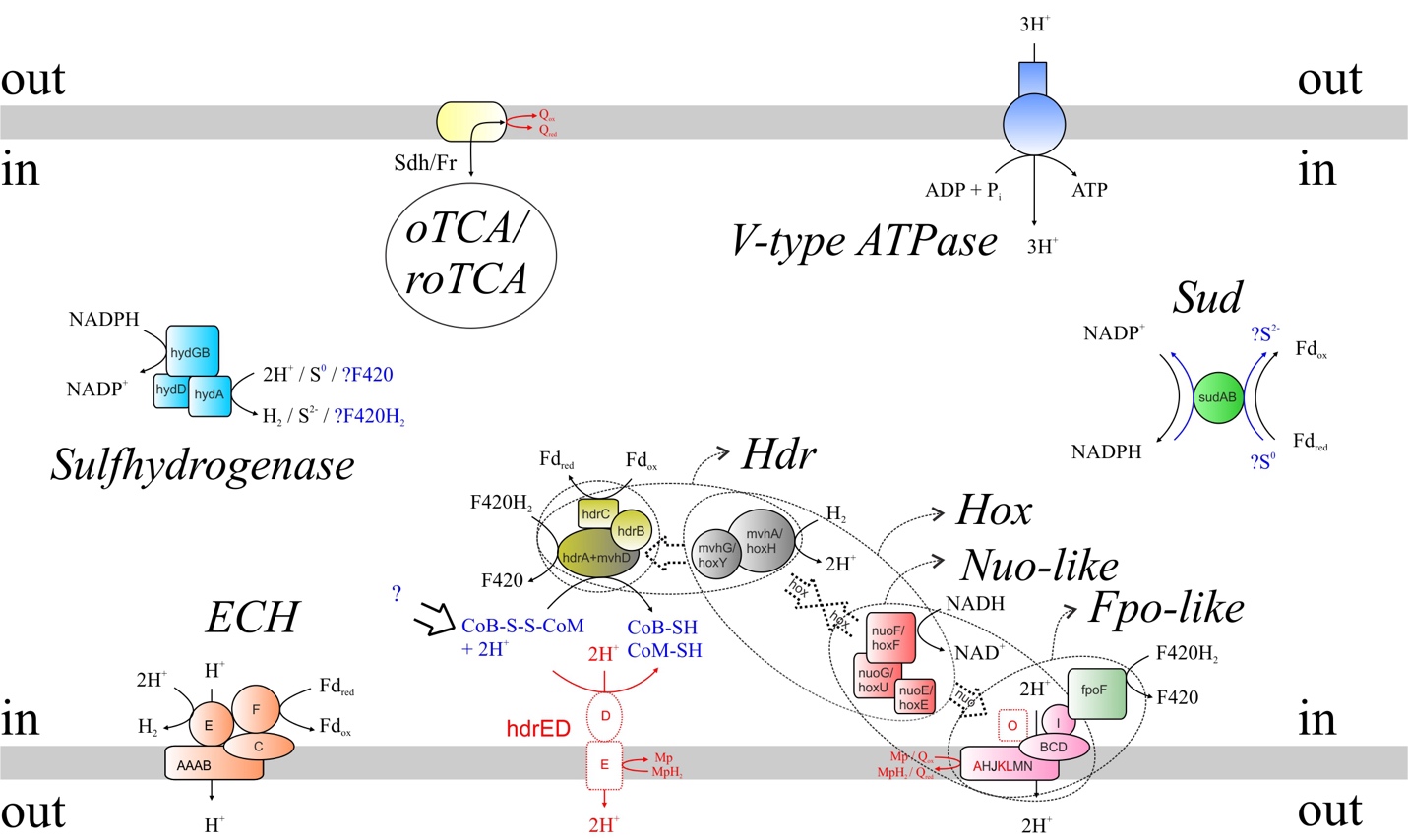

**Figure S2.** **Overview of main redox reactions and energy conservation in the cMAG; Bathy6-F90_cMAG_MillC.** All the enzyme complexes except ATP-synthase are shown with subunit detail. Protein abbreviations refer to the following gene ids in Table S1 (locus id from metaerg): ferredoxin: NAD^+^ oxidoreductase (RNF) is incomplete, only subunits C (1914) and D (1915) were found. Energy-converting hydrogenase (ECH) subunits C (551), E (552), B (553), F (554), A (557, 558, 559). Succinate dehydrogenase (Sdh, 371,372). Proton translocating V-type ATP-synthase: (15 – 22, 24). Six dehydrogenases/hydrogenase complexes were indicated as 1-6. 1: Putative F420H2-reducing, Fd-reducing hydrogenase/heterodisulfide reductase Mvh/Hdr: MvhA (2706, a NiFe family 3 hydrogenase), MvhG (2707), HdrA+MvhD (2887, a fused protein), HdrB (2745), HdrC (2744). 2: Hydrogenase Hox: HoxH (2706), HoxY (2707), HoxU (2708), HoxF (2709), HoxE (2710). 3: NADH dehydrogenase Nuo (Complex I) : NuoG (2708, a FeFe hydrogenase), NuoF (2709), NuoE (2710), NuoB (631), NuoCD (632, a fused protein), NuoN (633), NuoH (635), NuoL (636), NuoI (374). 4: F420H2 dehydrogenases Fpo: FpoF (2177), FpoJ (591), FpoB (631), FpoCD (632, a fused protein), FpoN (633), FpoH (635), FpoM (636), FpoI (374). 5: Sulfhydrogenase – NiFe (family 3) hydrogenase Hyd / sulfhydrogenase (Ma et al., 1993): HydGB (2711, a fused protein), HydD (2712), HydA (2713, a NiFe family 3 hydrogenase). This enzyme might reduce protons and elemental sulfur by NADH. 6: Sulfide dehydrogenase / Fd:NADH oxidoreductase Sud (Ma et al., 2001): SudA (671), SudB (672) which most probably involved in Fd-dependent NAD(P)H reduction or possibly can reduce sulfur with NADH. The genome also encoded a homolog (possibly a paralog) of SudA (2736) but no detectable SudB homolog. Complexes 1 - 4 represent alternative enzyme configurations since they possess common subunits with similar probability of function prediction. Despite being in the same genetic cluster HoxH/MvhA, HoxY/MvhG, HoxU/NuoG, HoxF/NuoF, HoxE/NuoE (2706-2710) and HydGB, HydD, HydA (2711-2713) are located on different DNA strands. HoxH/MvhA (2706) and HydA (2713) are homologs and HoxY/MvhG (2707) and HydD (2712) are homologs. Each enzyme complex has its own color. Red text and lines represent missing subunits and reactions. Blue text and lines are reactions in question.

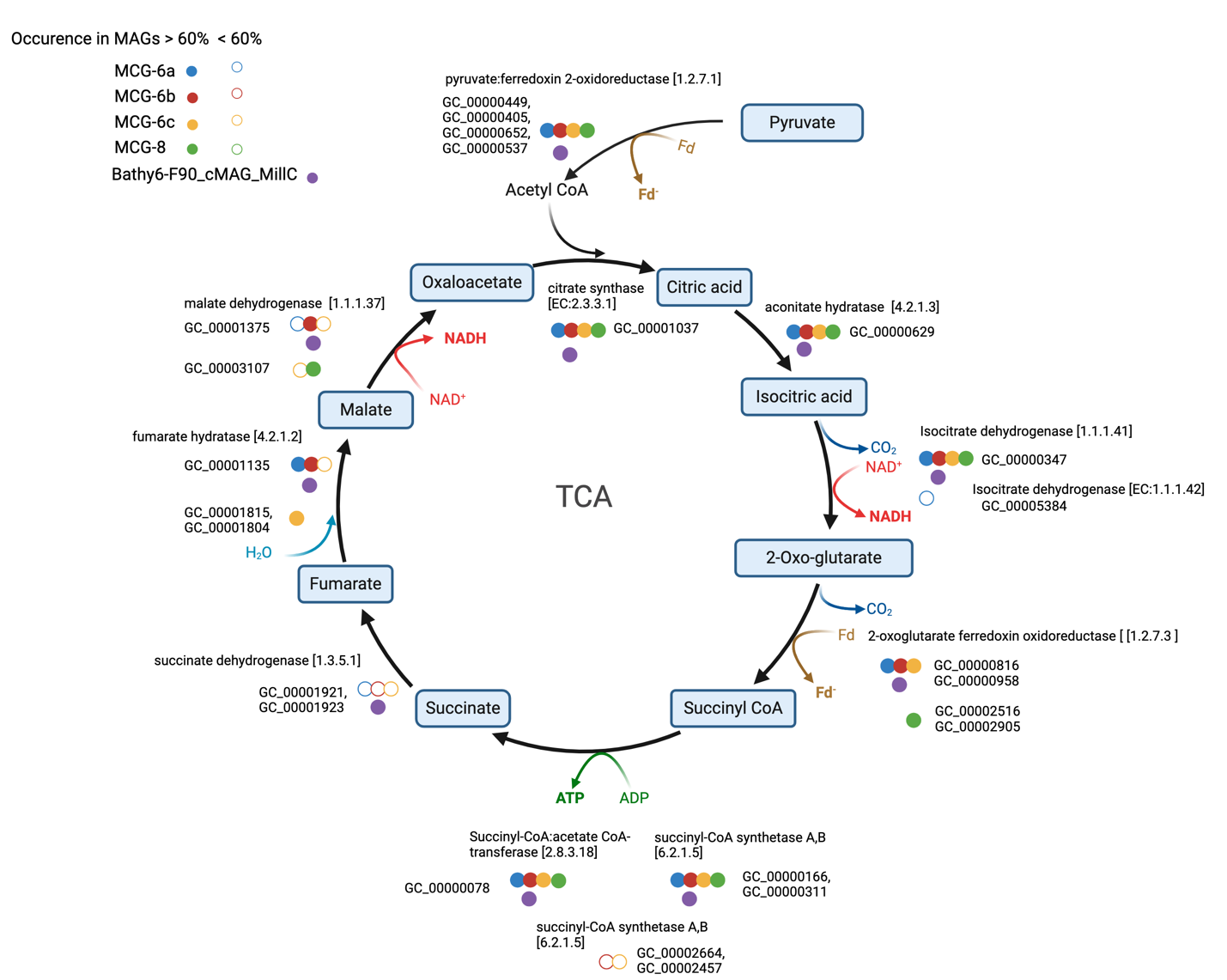

**Figure S3.**  Schematic of the TCA cycle in Bathy-6. Gene cluster IDs in the pangenome for the genes encoding each enzyme are given for each protein (Supplementary Table S3 and Table S4). The figure was drawn in Biorender.

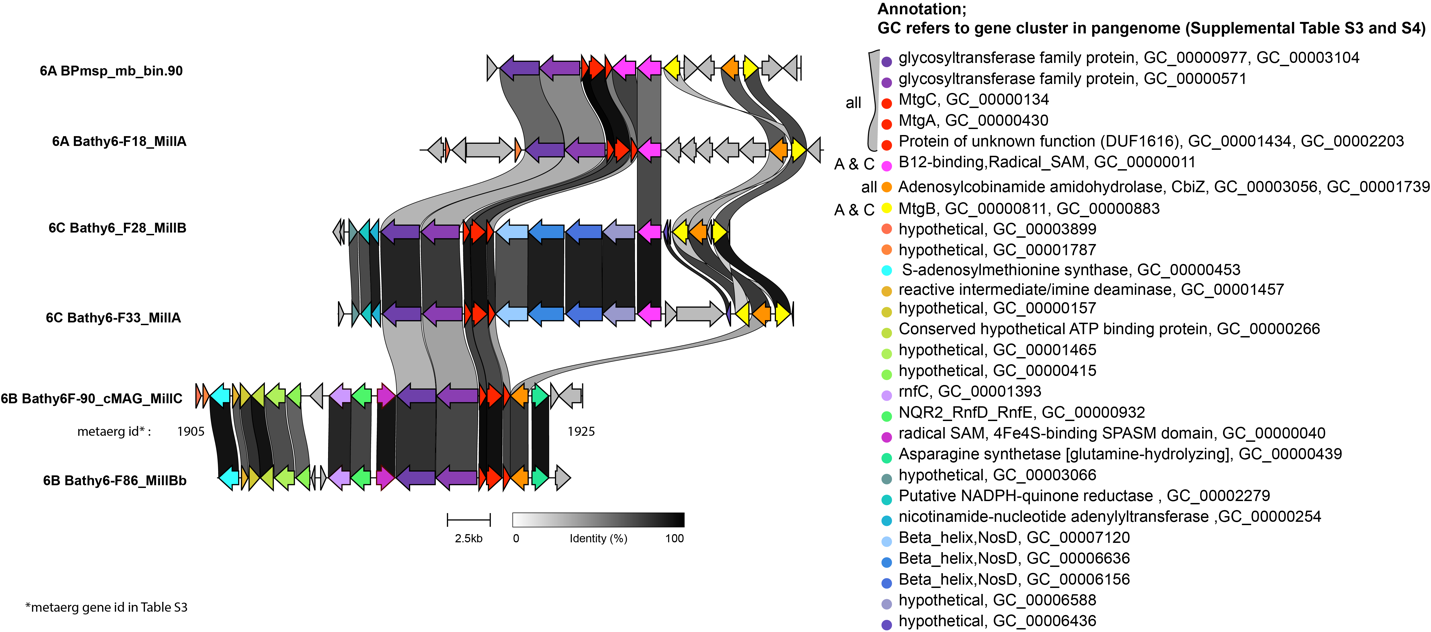

**Figure S4.** **Comparison of the genomic region containing the MtgA (a tetrahydromethanopterin S-methyltransferase subunit H (MtrH) homolog) and MtgC (a methanogenic corrinoid protein (MtbC1) homolog) genes encoding putative MT2 and MCP proteins.** Two MAGs from each Bathy-6 clade were included. Genes with a homolog in the other contigs or genomic regions are colored. The gene cluster id (GC_xxxxxxxx; Supplementary Table S3 and Table S4) and annotation are listed for these shared genes. All the Bathy-6 MAGs had the conserved gene cluster with the MtgA and MtgC genes (GC_00000430, GC_00000134), a hypothetical protein (GC_00001434, GC_00002203) and two glycosyl transferase family genes (GC_00000977, GC_00003104, GC_00000571). Homologs of the putative MtgB genes encoding the proposed substrate interacting o-demethylase (MT1) was only found in the Bathy-6 A and C clades (see Figure 1 for phylogenetic tree). Two of the GCs containing MtgB genes were found close to MtgA and MtgC (GC_00000811, GC_00000883). The third MtgB GC was always found unlinked to MtgA and MtgC (GC_00000053).

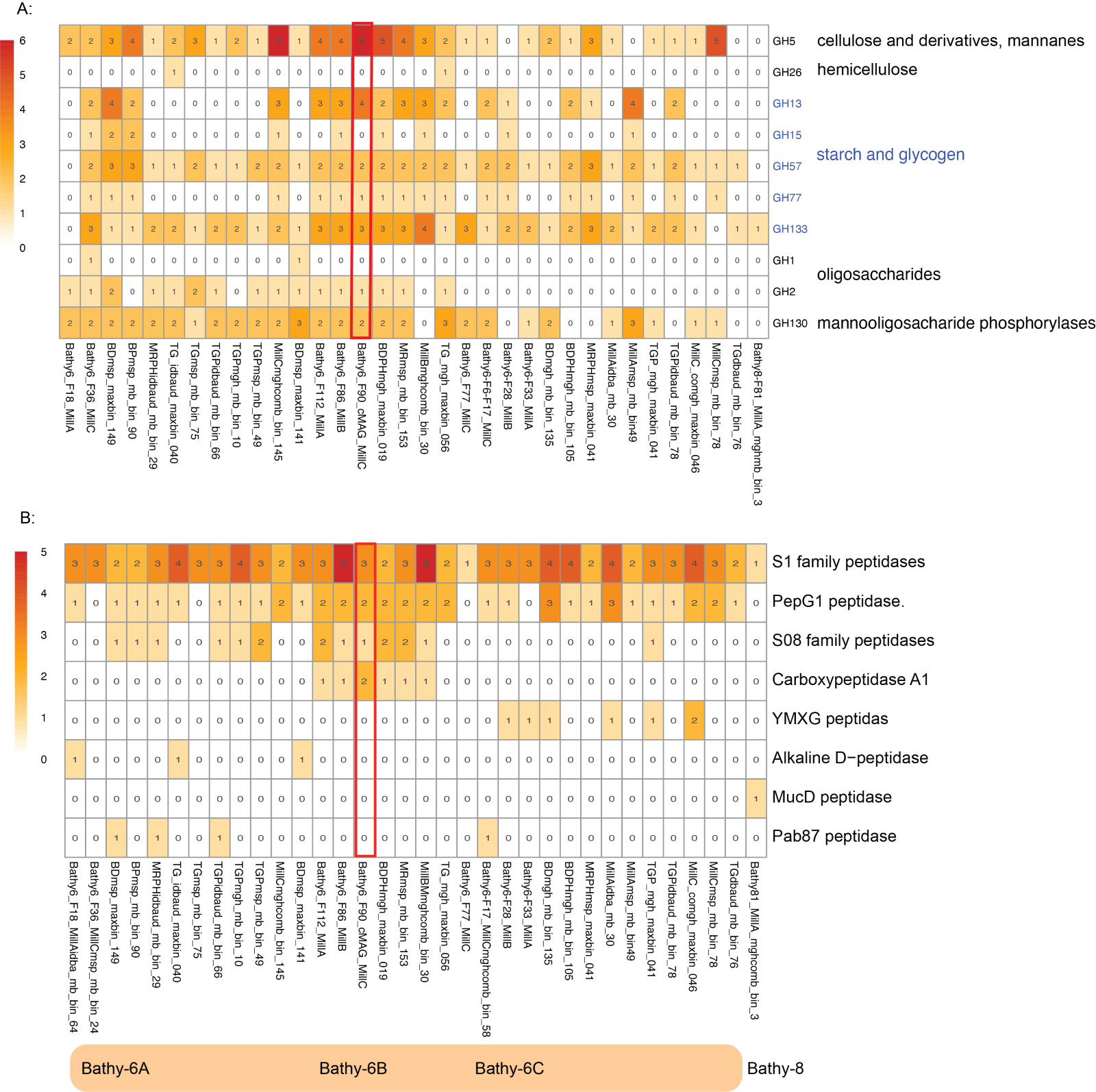

**Figure S5.** **Heatmap showing the distribution of gene clusters (GC_xxxxxxxx) annotated as CAZymes and peptidases in the MAGs constructed by us.** The MAGs analysed are on the x-axis and number of genes in each of the gene clusters (GC_xxxxxxxx; Supplementary Table S3 and Table S4) are shown as a heatmap along the y-axis. For the column showing the cMAG, Bathy6-F90_cMAG_MillC is outlined in red. Bathy-6A, 6B, and 6C refer to the Bathy-6 sub-clades in Figure 1. The Figure was drawn using pheatmap 1.0.12 in R.

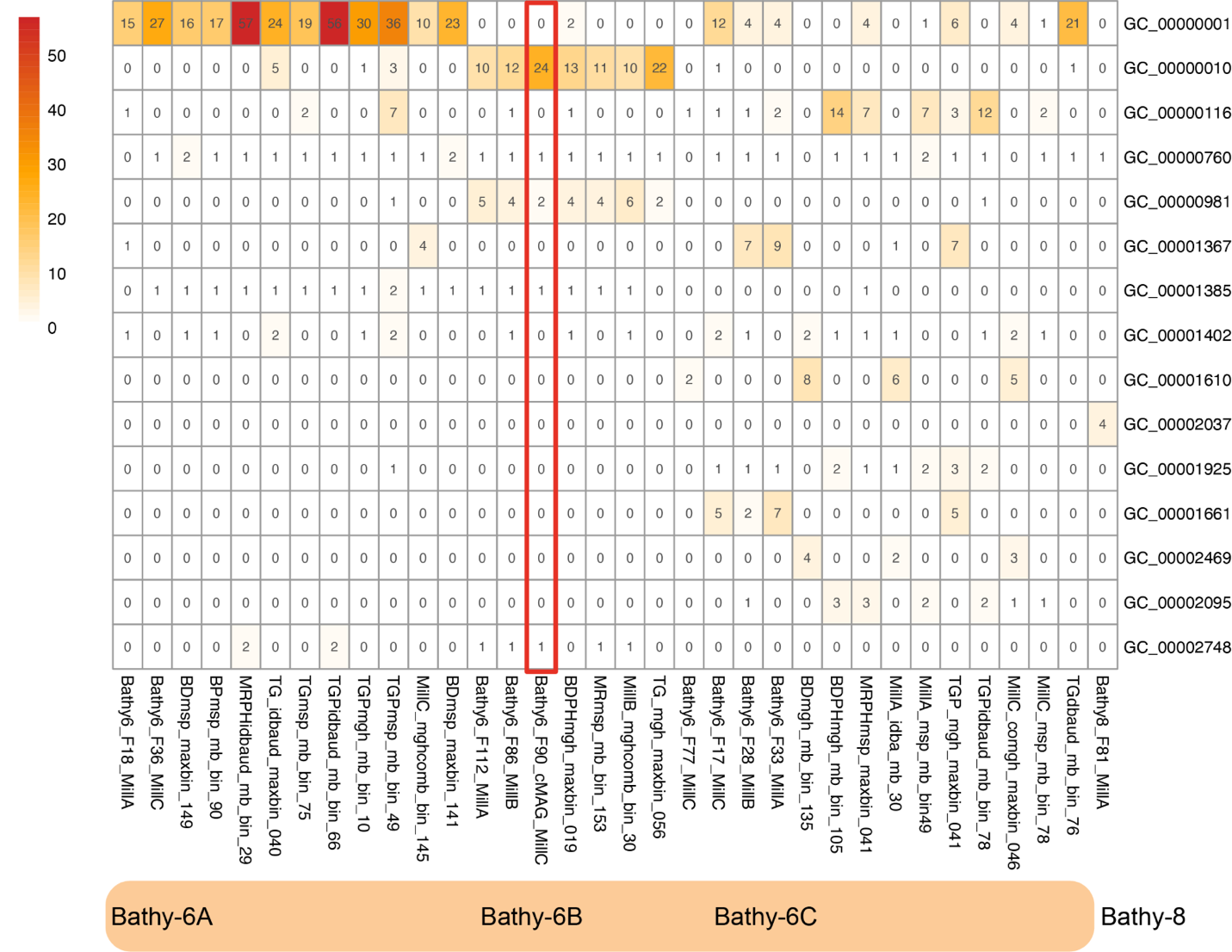

**Figure S6.** **Heatmap showing the distribution of the 15 most abundant PQQ-domain encoding gene clusters in the MAGs constructed by us.** The MAGs analysed are on the x-axis and number of genes in each of the gene clusters (GC_xxxxxxxx; Supplementary Table S3 and Table S4) are shown as a heatmap along the y-axis. The column showing the cMAG, Bathy6-F90_cMAG_MillC, is outline in red. Bathy-6A, 6B and 6C refers to the Bathy-6 sub-clades in Figure 1. The Figure was drawn using pheatmap 1.0.12 in R.

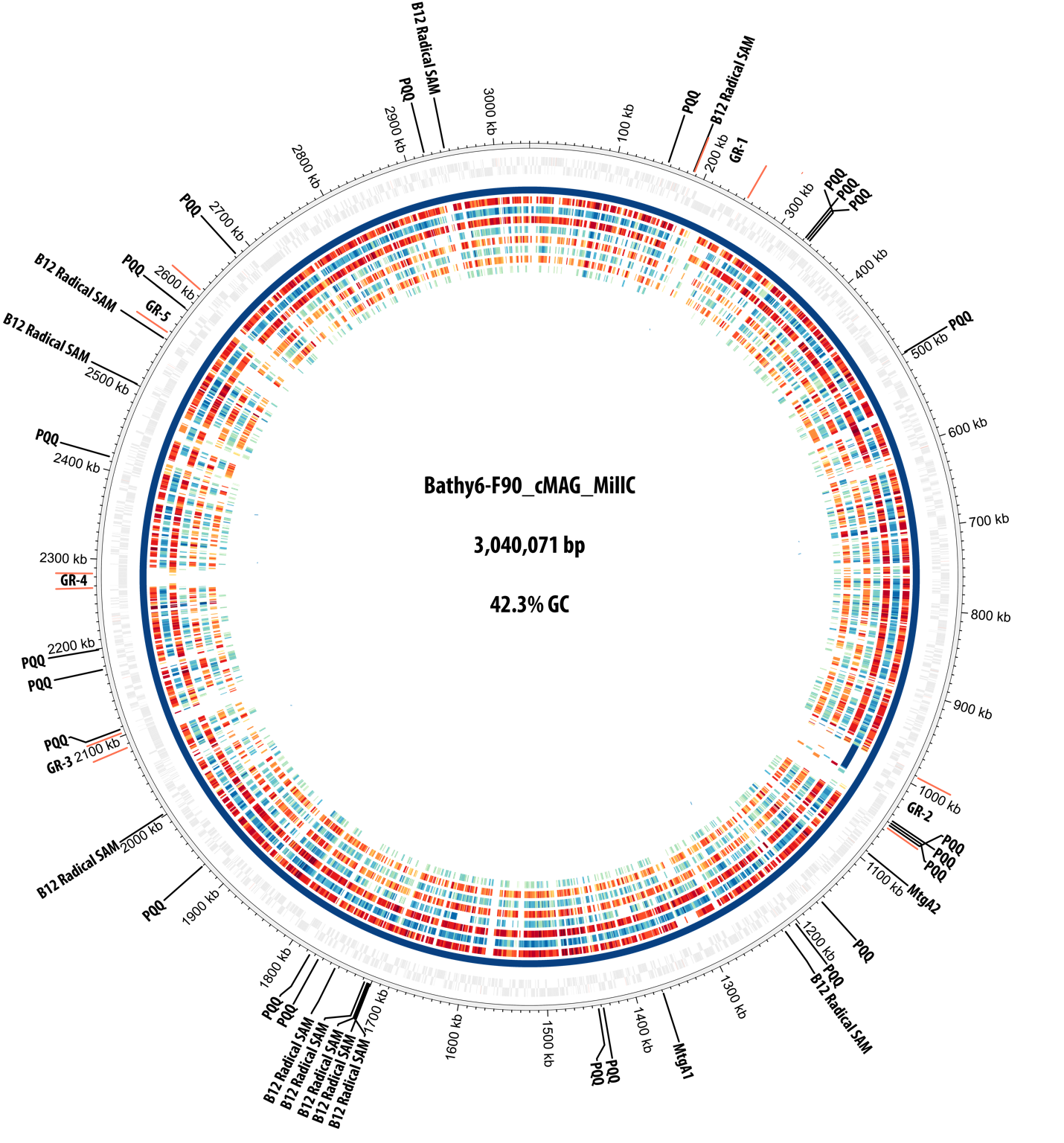

**Figure S7. Comparison of the Bathy6-F90_cMAG_MillC to the other high-quality MAGs assembled.** From outer to inner the circles represent: 1 numbering of the Bathy6-F90_cMAG_MillC genome indicating position of genomic regions discussed in the text (GR1-5), genes from GC_00000010 encoding proteins with PQQ-domains, genes encoding Radical SAM proteins with B12-binding domain with COG annotation COG1032, location of the gene cluster with the MtgA1 (and MtgC), location of the second MtgA gene (MtgA2). 2,3: CDS in the Bathy6-F90_cMAG_MillC genome on + and – strand respectively, 4: all Bathy6-F90_cMAG_MillC genes, gene homologous to Bathy6-F90_cMAG_MillC in  5: Bathy6-F86-MillB, 6: BDPHmgh_maxbin.019, 7: Bathy6_F112_MillA, 8: Bathy6_F18_MillA, 9: Bathy6_F36_MillC, 10: Bathy6-F28_MillB, 11: BDPHmgh_mb_bin105, 12: Bathy6_F81_MillA.

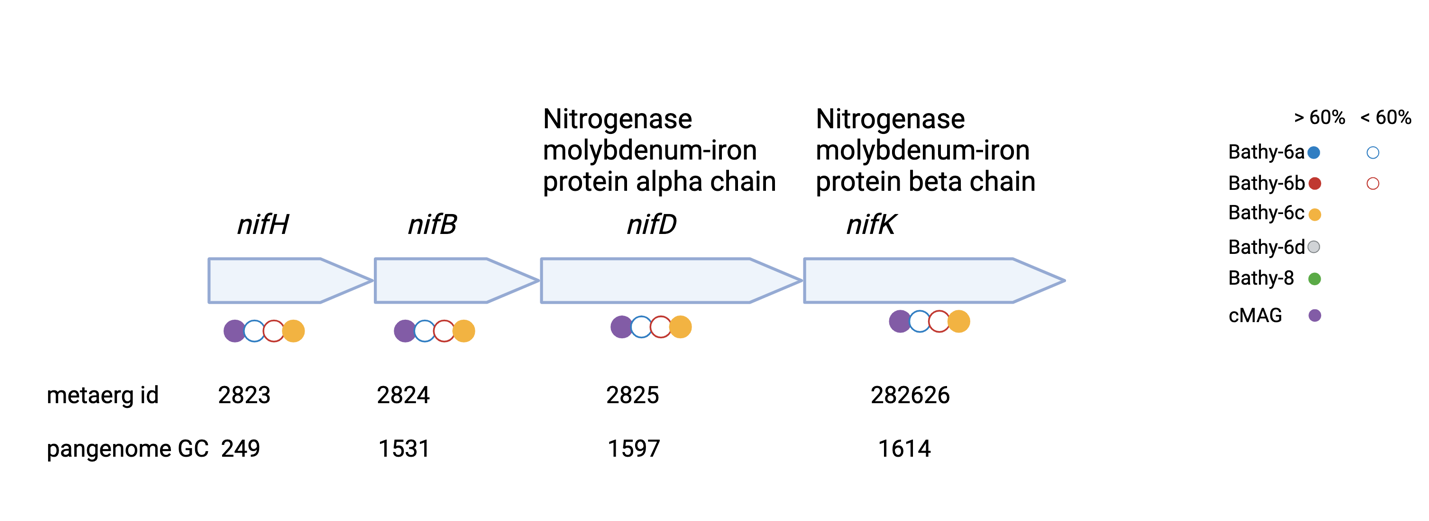

**Figure S8. Nitrogenase gene cluster.** Schematic of the nitrogenase gene cluster identified in the Bathy6-F90_cMAG_MillC genome and many Bathy-6 MAGs. Metaerg-id referres to the gene id assigned by the metaerg annotation software and is reported in Table S3. Pangenome GC referes to the pangenome gene cluster the gene was assigned to and is reported in Table S4. The circles under each gene indicates the Bathy-6 lineage (see Figure 1) the MAGs that carry the gene belongs to.

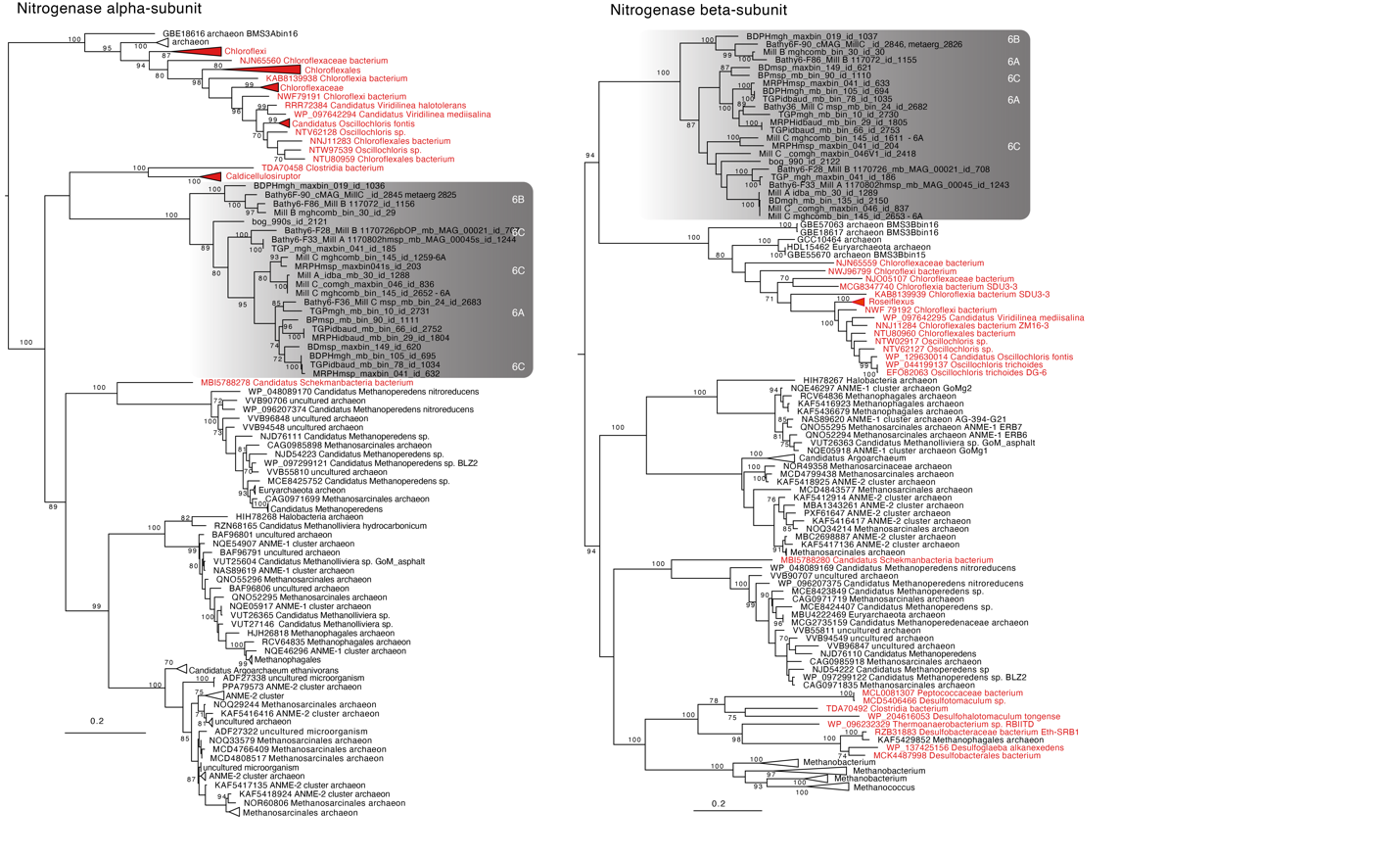

**Figure S9.** **Nitrogenase phylogeny.** Maximum likelihood phylogenetic trees of the alpha (panel a) and beta (panel b) subunits of nitrogenase enzymes. The 100 top blast matches obtained using the homolog from the Bathy6-F90_cMAG_MillC genome were downloaded from NCBI’s nr database and aligned to with the proteins from our MAGs using the MAFFT v 7.49 aligner in Geneious Prime 2022. Sites with > 90% gaps were removed, and the alignment was then used to construct a phylogenetic tree using RAxML v. 8.2.11 with the WAG + GAMMA model and 100 bootstrap replicates. Bootstrap support values >= 70 are shown on the branches. The shaded box indicates the Bathyarchaeia sequences identified in out MAGs and 6A, 6B and 6C next to the subclades refers to their lineage in the phylogenomic tree in Figure 1. Sequences labelled in red are Bacteria.

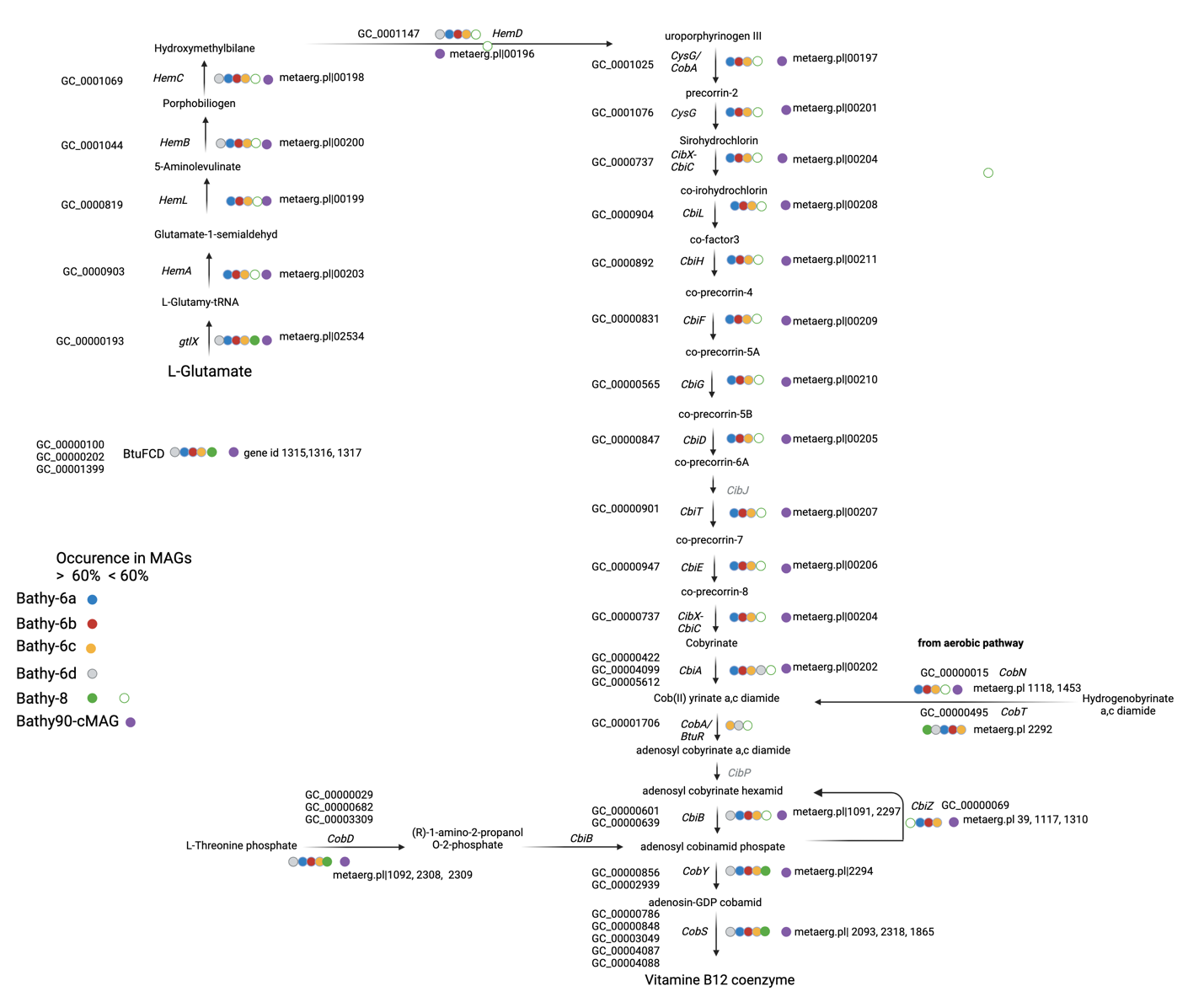

**Figure S10.** **Schematic of the vitamin B12 synthesis pathway inferred for Bathyarchaeia.**  For each enzyme the gene cluster id(s) (GC_xxxxxxxx) in the pangenome and metaerg id from the annotation of the closed genome, Bathy6-F90_cMAG_MillC, are indicated (Supplementary Table S3 and Table S4). The circles next to each enzyme indicates their presence in the genomes included in the pangenome analysis: Bathy6-F90_cMAG_MillC (purple circle), one of the Bathy-6 sub-clades or the Bathy-8 clade.

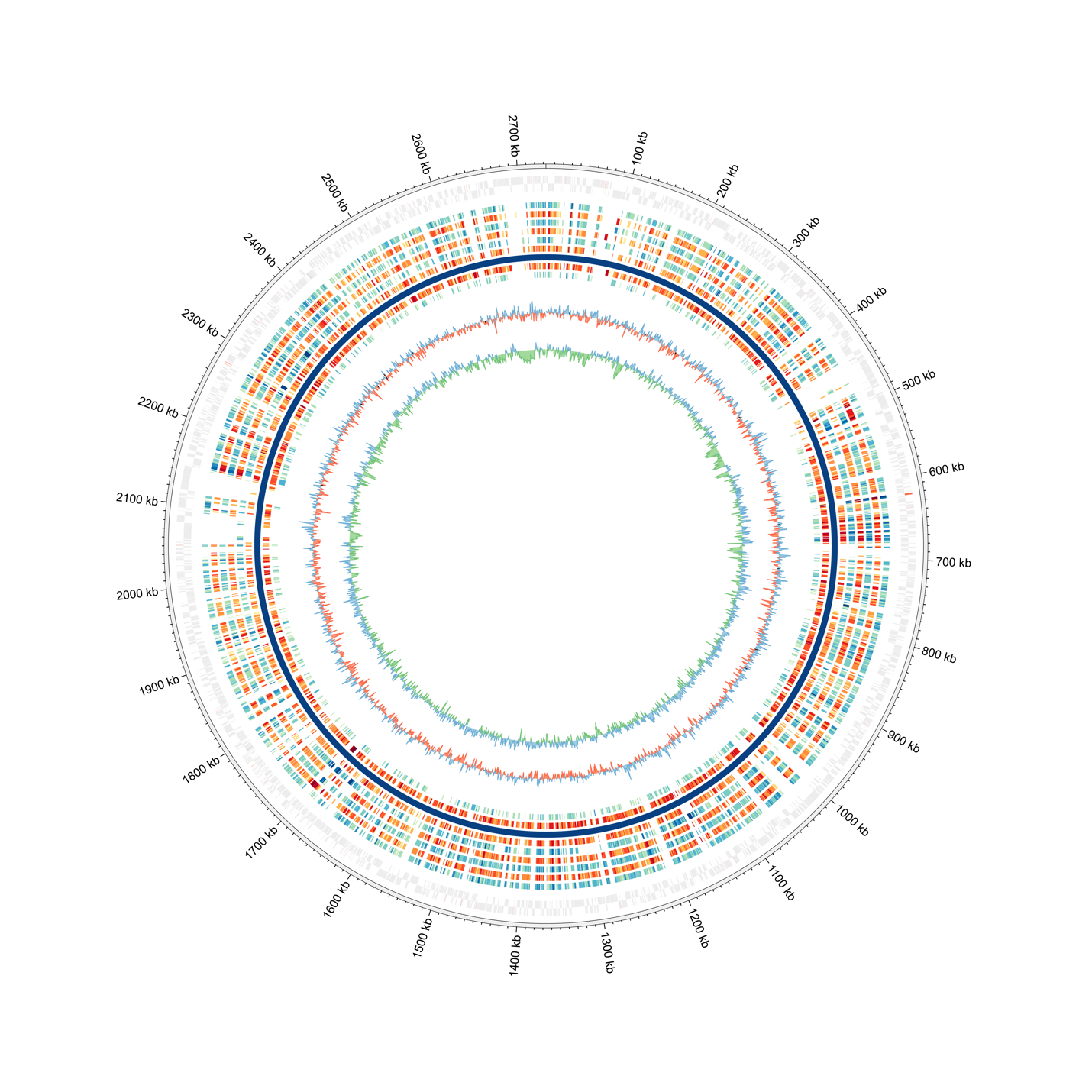

**Figure S11.** **Comparison of the Bathy6-F28_MillB to the other high-quality MAGs assembled.** From outermost to innermost the circles represent: 1; the Bathy6-F28_MillB MAG 2, 3: CDS in the Bathy6-F28_MillB genome on + and – strand respectively, gene homologous Bathy6-F28_MillB in  4: Bathy6-F90_cMAG_MillC 5: Bathy6-F86-MillB, 6: BDPHmgh_maxbin.019, 7: Bathy6_F112_MillA, 8: Bathy6_F18_MillA, 9: Bathy6_F36_MillC, 10: Bathy6-F28_MillB (self), 11: BDPHmgh_mb_bin105, 12: Bathy6_F81_MillA.

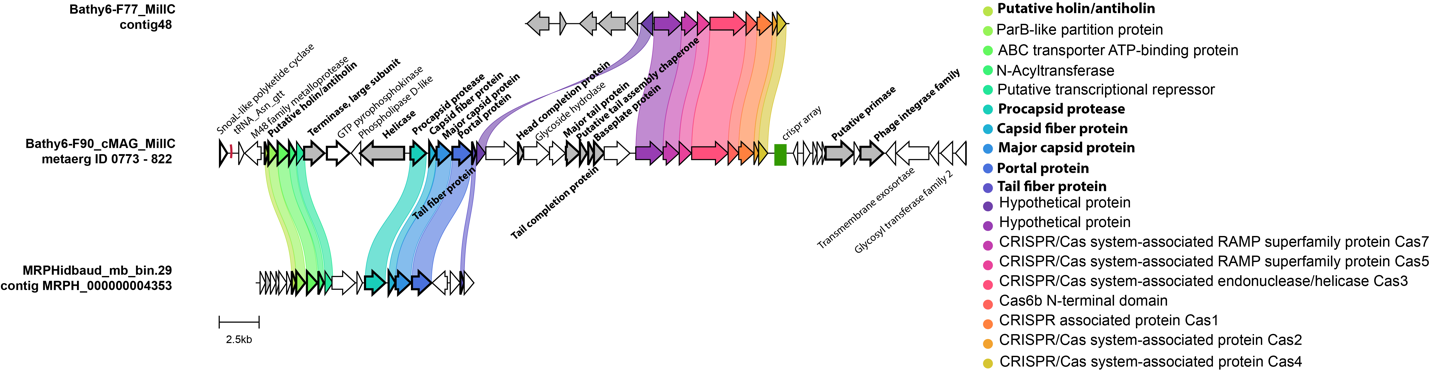

**Figure S12.** **Comparison of the putative provirus in Bathy6-F90_cMAG_MillC to contigs found in two of the other MAGs assembled here.** Typical virus genes are shown in bold. The figure was drawn using Clinker [34]. Interestingly the provirus contained a type I CRISPR-system [35] and a CRISPR repeat region. No Cas8 (or Cas10) gene were detected. Such streamlined elements are common in bacterioviruses [36].
